# The DC1 domain protein BINUCLEATE POLLEN is required for pollen development in *Arabidopsis thaliana*

**DOI:** 10.1101/2022.03.17.484815

**Authors:** Leonardo A. Arias, Sebastián D’Ippolito, Jésica Frik, Natalia L. Amigo, Fernanda Marchetti, Claudia A. Casalongué, Gabriela C. Pagnussat, Diego F. Fiol

**Author notes:** these authors contributed equally to this work (Leonardo A. Arias, Sebastián D’Ippolito, Jésica Frik). **Corresponding author**, D.F. Fiol. Instituto de investigaciones Biológicas IIB-CONICET- Universidad Nacional de Mar del Plata, Funes 3250, Mar del Plata, Buenos Aires, 7600, Argentina.

## Abstract

Development of the male gametophyte is a tightly regulated process that requires precise control of cell division and gene expression. A relevant aspect to understand the events underlying pollen development regulation constitutes the identification and characterization of the genes required for this process. In this work we showed that the DC1 domain protein BINUCLEATE POLLEN (BNP) is essential for pollen development and germination. Pollen grains carrying the defective *BNP* allele failed to complete mitosis II and are impaired in pollen germination. By yeast two-hybrid analysis and bimolecular fluorescence complementation assays, we identified a set of BNP-interacting proteins. Among confirmed interactors we found NAC family transcriptional regulators Vascular Plant One-Zinc Finger 1 (VOZ1) and VOZ2. VOZ1 localization changes during pollen development, moving to the vegetative nucleus at the tricellular stage. We observed that this relocalization requires BNP, as in the absence of BNP in pollen from *bnp/BNP* plants, VOZ1 nuclear localization is impaired. As *voz1voz2* double mutants showed the same developmental defect observed in *bnp* pollen grains, we propose that BNP requirement to complete microgametogenesis could be linked to its interaction with VOZ1/2 proteins. BNP could have a role of scaffold protein, recruiting VOZ1/2 to the endosomal system into assemblies that are required for their further translocation to the nucleus, where they act as transcriptional regulators.

## 1. Introduction

The development of the male gametophyte of flowering plants (pollen grain) takes place inside the anthers and involves two sequential phases: microsporogenesis and microgametogenesis. Microsporogenesis begins when a diploid microsporocyte undergoes meiotic division to form a tetrad of haploid microspores (Twell, 2011). During microgametogenesis a large vacuole is formed inside the released microspores and positions the nucleus in the periphery of the cell. This stage is followed by an asymmetric division or Pollen Mitosis I (PMI) that yields a large vegetative cell and a small germ cell. The germ cell is subsequently engulfed within the cytoplasm of the vegetative cell. The germ cell then undergoes a second division, Pollen Mitosis II (PMII), which results in twin sperm cells and yields the mature tricellular pollen. In species such as *Arabidopsis thaliana,* PMII takes place inside the pollen grain prior to anthesis. Although transcriptional studies have provide significant advances in the understanding of the molecular mechanisms controlling male gametophyte development, the picture is still far from complete (reviewed in (Hafidh et al., 2016). Studies on isolated mutants have also proved to be of great importance to identify novel genes and cellular mechanisms involved in this process (Borg et al., 2009).

Divergent C1 (DC1) domains are cysteine/histidine-rich zinc finger modules found exclusively in the plant kingdom. DC1 domains resemble the C1 domains present in protein kinase C (PKC) and in over a dozen of animal proteins (Colon-Gonzalez and Kazanietz, 2006). However, C1 domains are not frequent in plants, as they were only identified in diacylglycerol kinases (Escobar-Sepúlveda et al., 2017). C1 domains have the ability to bind the secondary messenger diacylglycerol (DAG) but also participate in protein-protein interactions and in the targeting of proteins to the membrane (Colon-Gonzalez and Kazanietz, 2006). Sequence and structure similarities between DC1 and C1 domains suggest that some of the roles played by C1 domains in animal proteins could be fulfilled in plants by DC1 domains (D’Ippólito et al., 2017).

The Arabidopsis genome encodes for over 140 uncharacterized proteins harboring DC1 domains (D’Ippólito et al., 2017). Reports regarding genes coding for DC1 domain proteins were limited to transcriptional responses. In Arabidopsis, *At5g17960* was described as responsive to hormones and stress treatments (Bhaskar et al., 2015) and *ULI3 (At5g59920)* was related to UV-B responses (Suesslin and Frohnmeyer, 2003). In other plant species, DC1 domain proteins have been also associated with stress and hormone responses (Shinya et al., 2007; Li et al., 2010; Hwang et al., 2014; Gao et al., 2016).

The first functional characterization of a DC1 domain protein, Vacuoleless Gametophytes (VLG), was recently reported (D’Ippólito et al., 2017). VLG localizes to prevacuolar compartments/multivesicular bodies and it was shown to be essential for the development of the female and male gametophytes. A role for VLG was proposed during vacuole biogenesis, likely through the interaction with the VAMP protein PVA12 and the GDSL-motif lipase LTL1 (D’Ippólito et al., 2017).

In this work we studied one of the closest VLG homologs, encoded by *At2g44370* and identified that the DC1 domain protein BINUCLEATE POLLEN (BNP) plays a relevant role during the male gametophyte development in Arabidopsis. In addition, we demonstrated that BNP is required for pollen germination and early sporophytic developmental stages. To functionally characterize BNP we searched for protein interactors and identified transcriptional regulators VOZ1 and VOZ2 as BNP binding proteins and we showed that VOZ proteins co-localize to BNP, supporting a physiological interaction. We showed that VOZ1 localizes in the cytoplasm at early stages of pollen development, but moves to the vegetative nucleus at the tricellular stage. We observed that this relocalization requires BNP, as in the absence of BNP in pollen from *bnp/BNP* plants, VOZ1 nuclear localization is impaired. As *voz1voz2* mutants also exhibit pollen developmental defects (Celesnik et al., 2013), we propose that the interaction between BNP and VOZ proteins outside the nucleus might impact further on VOZ1/2 activities, resulting in functional consequences for gene expression. Thus, BNP could act as a scaffold protein recruiting VOZ1/2 and likely other proteins to the endosomal system into assemblies that might be required to accomplish VOZ1/2 translocation to the nucleus.

## 2. Results

### BNP codes for a DC1 domain containing protein

*At2g44370,* here named *BINUCLEATE POLLEN (BNP),* is a single exon gene that encodes for a 250-amino acid long protein with three DC1 domains (Fig. 1B). Phylogenetic analysis of Arabidopsis DC1 domain protein family grouped BNP into the most divergent cluster, with At5g40590 and VLG being the closest homologs to BNP (Fig. 1C). Multiple sequence alignments of selected protein sequences of this cluster and DC1 domain proteins from *Capsicum annuum,* tobacco, wheat and cotton showed amino acid conservation mostly restricted to the residues that define the DC1 signature motif, which might be involved in Zn^2+^ coordination and folding of the domain (Fig. 1D). On the other hand, amino acids corresponding to loop regions in the folded domain - which might define its binding capabilities (Rahman and Das, 2015)- showed very low similarity, suggesting a diversity of interactors for the aligned proteins.

**Figure 1.**
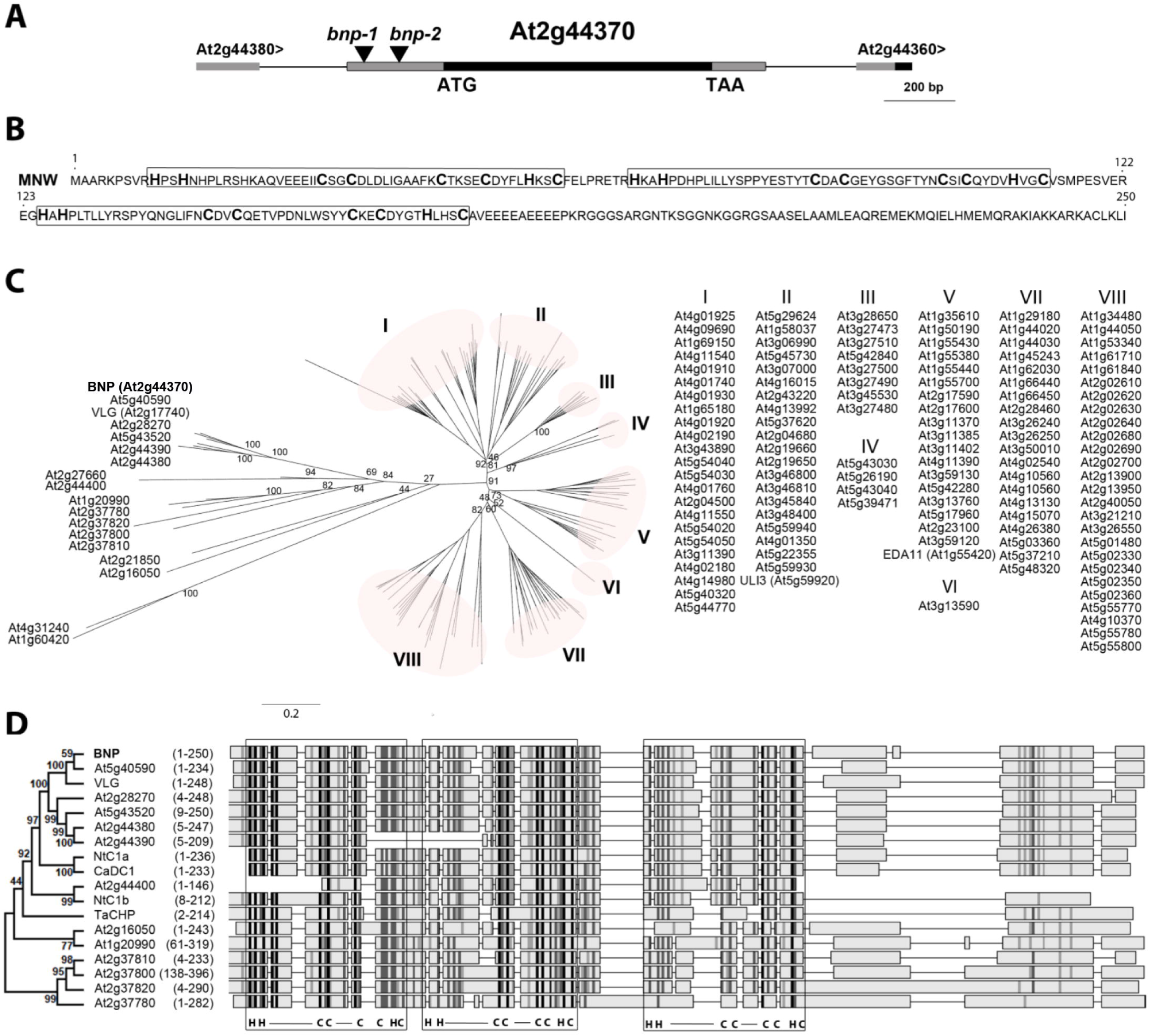
BNP codes for a DC1 domain containing protein. (A) Schematic view of a region of Arabidopsis chromosome 2 (from 18,323,538 to 18,321,537 bp -TAIR10) showing BNP genomic location and bnp-1 and bnp-2 insertion sites. (B) BNP amino acid sequence with boxed DC1 domains. (C) Phylogenic tree of 140 DC1 domain containing proteins in Arabidopsis. Bootstrap test results (1000 replicates) of the major nodes are indicated. IDs of sequences grouped in clusters I to VIII are listed. (D) Alignment of BNP with its closest Arabidopsis homologs and CaDC1 (AEI52549), TaCHP (ACU80555), NtDC1a (BAF80452) and NtDC1b (BAF80453). Phylogenic tree with bootstrap test results (1000 replicates) for the alignment of the complete 18 sequences is shown. Amino acid region represented for each sequence is indicated. DC1 domains are framed, signature residues and binding loop regions are indicated (C=Cys, H=His, — = loop). Higher intensity in grey scale denotes higher sequence similarity.

### *bnp* is a gametophytic-acting mutation

Two Arabidopsis lines with T-DNA insertions in *BNP* (SALK_114889 –here named *bnp-1* and GK-008E01 -here named *bnp-2)* were studied (Fig. 1A). After backcrossing to the Col-0 ecotype, the progeny of self-pollinated plants was genotyped. No homozygous mutant plants were recovered in the offspring from *bnp-1/BNP* or *bnp-2/BNP* self-pollinated plants. The ratio of hemizygous to WT plants in the progeny of self-fertilized *bnp-1/BNP* plants was 1.44 (*n* = 183). For *bnp-2/BNP* plants, segregation analysis for sulfadiazine resistance showed a ratio of resistant to sensitive plants of 1.31 (*n* = 2634). The proportion of plants harboring the transgene recovered in the progeny of self-fertilized plants was lower than the Mendelian segregation ratio expected for diploid sporophytic (3:1) or embryo lethal (2:1) mutants suggesting a gametophytic defect. Furthermore, no obvious sporophytic phenotypes were observed in the hemizygous plants, suggesting that *bnp* is a recessive mutation (Fig. S1). The siliques of *bnp/BNP* plants showed a small percentage of aborted seeds, not different to WT plants (Fig. S1).

As no homozygous plants were recovered and siliques from hemizygous plants looked normal, the *BNP* mutation could be affecting pollen development, germination or early development. To test this hypothesis, we performed reciprocal crosses between *bnp/BNP* and WT plants and calculated the transmission efficiency of the mutant allele (TE = resistant/WT offspring x 100) (Table 1). Only 15.5% of microgametophytes carrying *bnp-1* and 22.0 % of the microgametophytes carrying *bnp-2* transmitted the insertion to the next generation (p<0.0001, Chi square test), indicating that *bnp* affects male gametophyte development or function (Table 1). No significant defects were observed for transmission through the female gametophyte (90.8% and 95.9% for *bnp-1* and *bnp-2,* respectively, Table 1, p-values 0.51 and 0.66 respectively, Chi-square test).

**Table 1.** Transmission efficiency of *bnp* alleles. Transmission efficiency of the *bnp-1* and *bnp-2* alleles in reciprocal crosses between mutant and WT plants.

| Parental genotypes |  | Genotype of progeny† |  | Transmission efficiency of: | p-value* |
| --- | --- | --- | --- | --- | --- |
| Female | Male | WT | *bnp-1/BNP* | *bnp-1* |  |
| *bnp-1/BNP* | WT | 98 | 89 | ♀ 90.8% | 0.51 |
| WT | *bnp-1/BNP* | 181 | 28 | ♂ 15.5% | <0.0001 |
|  |  | WT | *bnp-2/BNP* | *bnp-2* | p-value* |
| *bnp-2/BNP* | WT | 217 | 208 | ♀ 95.9% | 0.66 |
| WT | *bnp-2/BNP* | 254 | 56 | ♂ 22% | <0.0001 |

†No homozygous plants were detected; \* Chi-square test for a 1:1 segregation hypothesis (degrees of freedom = 1).

Although the insertions in *BNP* compromised the transmission of the mutation through the male gametophyte, a fraction (about 15-22%) of pollen grains are still be able to transmit the insertion to the next generation. However, we did not recover homozygous mutant plants in the self-progeny of hemizygous plants. Thus, we investigated whether embryogenesis or seed germination were affected by the mutation analyzing pistils and seeds from *bnp/BNP* plants. No obvious embryo defects were observed in siliques from *bnp/BNP* plants, as neither developing siliques nor seed sets showed differences from WT siliques (Fig. S1). However, seeds from *bnp/BNP* self-pollinated plants showed reduced germination rates when compared to WT plants (Table 2). Further analysis of the germinated seeds showed that a fraction of the seedlings arrested soon after germination. Values of seedling lethality obtained were of 0.9% for WT (*n* = 440), 3.4% (*n* = 754) for seeds from *bnp-1/BNP* self-pollinated plants and 4.5% (*n* = 973) for *bnp-2/BNP* self-pollinated plants (Table 2). PCR genotyping of arrested seedlings showed that 75% (*n* = 12) corresponded to *bnp/bnp* homozygous individuals.

**Figure 2.**
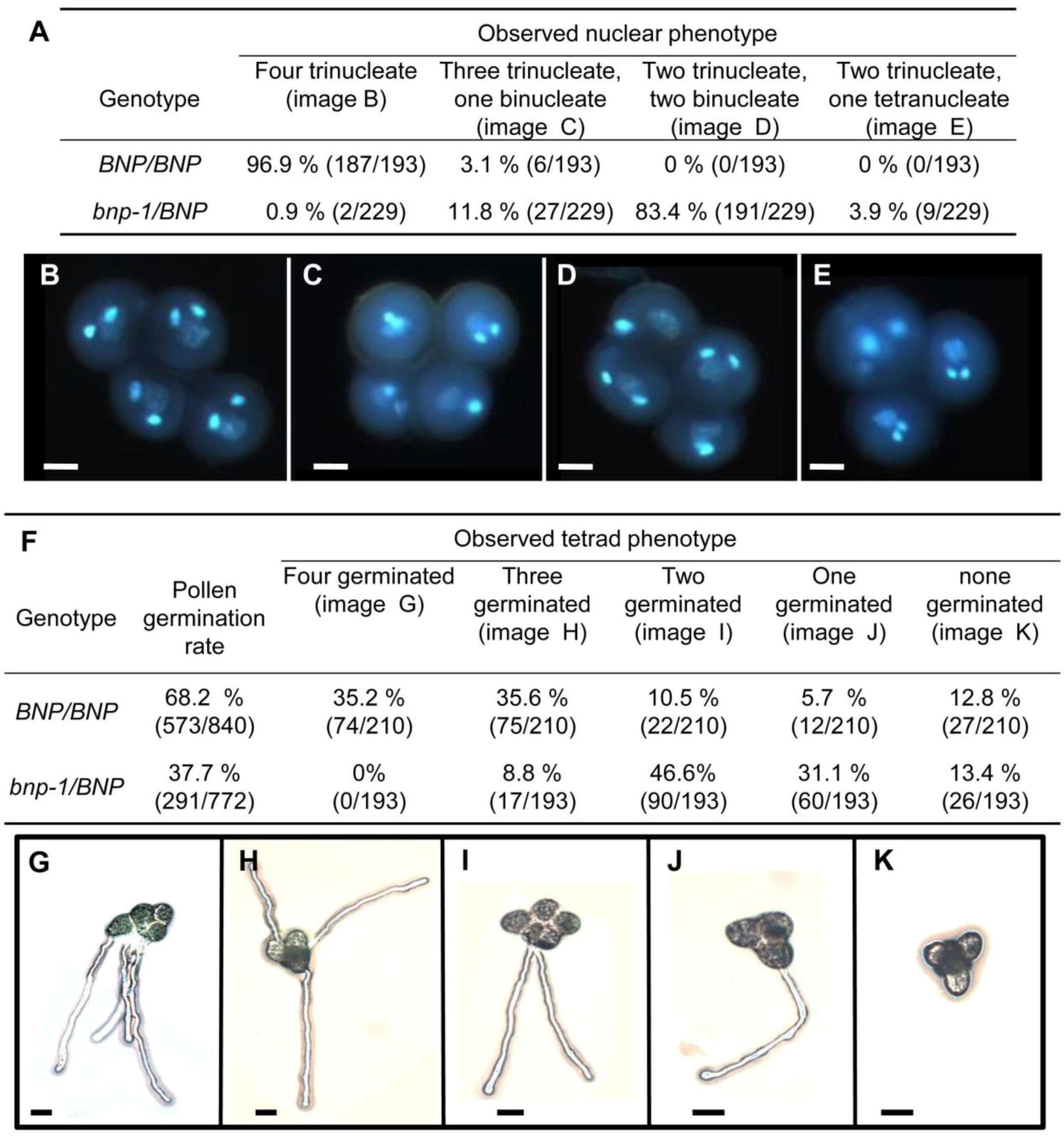
Pollen development and germination is impaired in *bnp-1* mutants. (A) Quantification of phenotypes observed in *BNP/BNP* and *bnp-1/BNP* plants in *qrt1/qrt1* background. (B-E) Nuclear configuration types revealed by DAPI (4’,6-Diamidino-2-phenylindole) staining. Bars = 10 μm. (F) Quantification of pollen tube germination in tetrads from *BNP/BNP* and *bnp-1/BNP* plants in *qrt1/qrt1* background. (G-K) Pollen tube germination in tetrads. Bars = 20 μm.

**Table 2.** Viability of the progeny in *bnp* mutants. Seed germination and seedling lethality of the progeny of self-pollinated *bnp-1/BNP, bnp-2/BNP* and WT plants.

| Parental genotype | % viable progeny | % unviable progeny |  | p-value* |
| --- | --- | --- | --- | --- |
|  |  | not germinated | seedling lethality |  |
| WT | 98.6 (435/441) | 0.2 (1/441) | 1.1 (5/441) |  |
| *bnp-1/BNP* | 94.8 (715/754) | 1.7 (13/754) | 3.4 (26/754) | <0.001 |
| *bnp-2/BNP* | 91.4 (688/753) | 2.8 (21/753) | 5.8 (44/753) | <0.00001 |

In parenthesis, number of individuals observed over the total analyzed. \* Chi-square test for a hypothesis of no relationship between the genotype of the parents and the viability of the progeny (degree of freedom = 1).

Altogether these results suggest that the mutation might cause impairment in male gametogenesis or function, reduced seed germination and seedling lethality, which combined, explain the absence of homozygous mutant plants in the progeny of selfed hemizygous plants.

### Pollen development in *bnp* mutants is arrested at the binuclear stage

Viability of mature pollen was analyzed in *bnp-1/BNP* and *bnp-2/BNP* by means of Alexander staining and FDA staining. No differences were observed compared to pollen obtained from WT plants, indicating that the *bnp* mutation does not affect pollen viability (Fig. S2).

We later analyzed nuclear composition of mature pollen using DAPI staining. Normal mature pollen contains three nuclei: a vegetative nucleus and two sperm nuclei generated by the mitotic division of the germ cell. Mature pollen from WT plants showed 99.2% (*n* = 772) of trinucleate grains. On the other hand, *bnp/BNP* plants presented only about 57-59% of trinucleate mature pollen grains (56.9% (*n* = 1123) in *bnp-1/BNP* and 59.0% (*n* = 981) in *bnp-2/BNP)* (Table 3). The rest were found in a binucleate stage, suggesting developmental arrest.

**Table 3.** Nuclear composition of mature pollen in *bnp* mutants. Mature pollen from *bnp-1/BNP, bnp-2/BNP* and WT plants was stained using DAPI.

| Genotype | % trinucleate pollen | p-value* |
| --- | --- | --- |
| WT | 99.2 (766/772) |  |
| *bnp-1/BNP* | 56.9 (639/1123) | <0.0001 |
| *bnp-2/BNP* | 59.0 (579/981) | <0.0001 |

In parenthesis, number of individuals observed over the total analyzed. \* The statistical significance was determined by Student's *t*-test.

A genetic complementation assay confirmed that the insertions in *BNP* are causing the abnormalities observed during pollen development. *bnp-1/BNP* hemizygous plants transformed with a *ProBNP:BNP:GFP* translational fusion construct showed trinucleate mature pollen grains at similar levels of WT plants (Fig. S3). This result was verified in T3 plants of five independent complemented lines.

In order to better characterize the defect observed during pollen development, *bnp-1/BNP* plants were crossed to *qrt1* homozygous plants, which are impaired in a pectin methylesterase responsible for the separation of microspores tetrads (Francis et al., 2006). F2 plants that were hemizygous for the insertion in *BNP* and homozygous for *qrt1 (bnp/BNP qrt1/qrt1)* were further analyzed. Mature tetrads were collected and stained with DAPI. Consistently with our previous results, most tetrads (99.1%) of *bnp-1/BNP qrt1/qrt1* plants contained at least one microspore and, in most cases, two microspores with an abnormal number of nuclei (Fig. 2A), indicating a failure to complete PMII in about 46% of the pollen grains analyzed.

We next assessed germination of pollen grains in the tetrads. As shown in Fig. 2, germination rates were 37.7% in pollen collected from *bnp-1/BNP qrt1/qrt1* plants and 68.2% in pollen grains from *BNP/BNP qrt1/qrt1* plants (Fig. 2F). This indicates a reduction of 44.7% in the germination rate in pollen from *bnp-1/BNP qrt1/qrt1* plants compared to pollen grain tetrads from *BNP/BNP qrt1/qrt1* plants. As the reduction in the rate of germination observed in the hemizygous background *bnp-1/BNP* was less than 50%, this suggests that although severely affected, a small fraction of *bnp* pollen grains are still able to germinate.

Analysis of pollen grain germination in *bnp-2/BNP* plants showed comparable values, as we calculated a decrease in germination rate of about 36% compared to pollen grains from WT plants. Germination rates of 43% (*n* = 787) and 67% (*n* = 443) were recorded for pollen grains collected from *bnp-2/BNP* and WT plants respectively.

These results showed that *BNP* is required to complete pollen development, specifically to undergo PMII. Arrested pollen grains were viable, but germination was impaired, which explains the reduced transmission of *bnp* through the male gametophyte (Table 1).

### *BNP* is expressed in pollen and young sporophytic tissues

Expression of *BNP* was studied in Arabidopsis pollen expressing the *BNP-GFP* fusion driven by the *BNP* promoter, which was proved to be functional, as it complemented the phenotype in *bnp/BNP* plants (Fig. S3). BNP-GFP was detected in cytoplasmic speckles following a punctuate pattern at early stages of pollen development (microspore tetrad and released microspore) and in discrete bigger compartments at bicellular and tricellular stages (Fig. 3A and 3H). The same localization pattern was observed in the pollen tube (PT) of germinating pollen grains (Fig. 3I-J).

**Figure 3.**
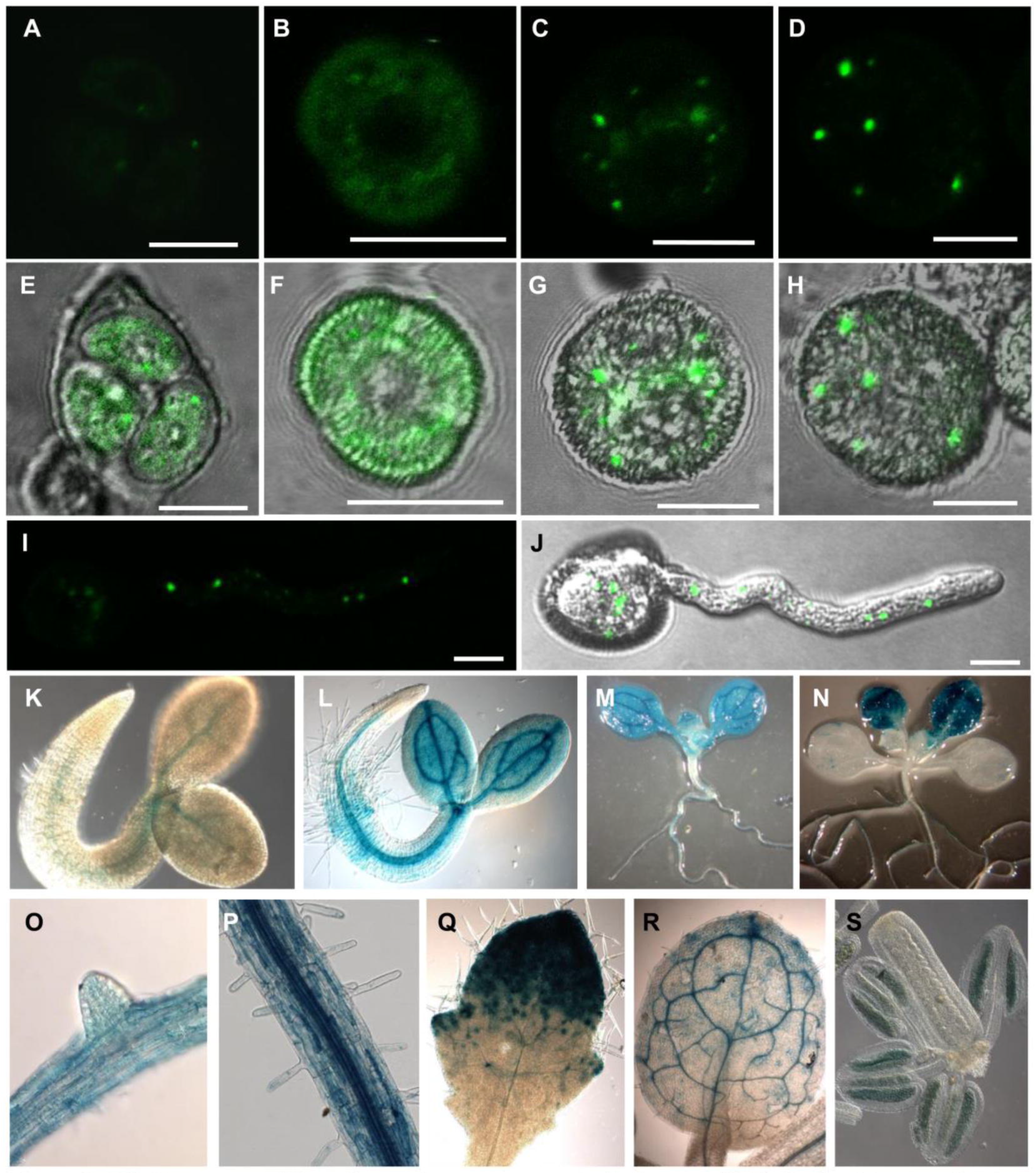
*BNP* is expressed in pollen and young sporophytic tissues. (A-D) Confocal images showing BNP-GFP through microgametogenesis. (E-H) Overlay between DIC and GFP fluorescence. (A and E) Microspore tetrad, (B and F) released microspore, (C and G) bicellular stage and (D and H) tricellular stage. (I) Image showing a projection along the z axis of all planes showing GFP fluorescence. (J) Overlay of DIC and GFP fluorescence showing a germinating pollen grain. (K-S) GUS activity in Arabidopsis transgenic plants carrying the *pBNP:GUS* construct. (K) Seedling 24 h after imbibition, (L) 48 h after imbibition, (M) 6 days post germination and (N) 12 days post germination. (O) Root with lateral root primordium, (P) primary root with root hairs, (Q) third leaf 8 days after imbibition (R) third leaf 14 days after imbibition. (S) GUS signal was detected inside anthers at floral developmental stage 10. Three independent transgenic lines were used for this study. Pictures are representative of the results obtained in all analyzed lines. Bars = 10 μm.

To gain further knowledge of BNP roles *in planta,* a reporter construction expressing the *GUS* gene under the control of *BNP* promoter was used to analyze *BNP* expression. GUS activity accumulated at early stages of seedling development, mainly in vascular tissues and developing leaves. It was also observed associated with anthers at very early stages of floral development, indicating that expression of *BNP* can be detected before meiosis (Fig. 3K-S). Expression data observed were comparable to transcriptional data available in the efp Browser Data Analysis Tool (https://bar.utoronto.ca/eplant/) (Fig. S4). These results also indicated that BNP could be functional not only during pollen development but also at additional developmental stages, such as sporogenesis, germination and early stages of sporophyte development.

### BNP interacts with transcription factors VOZ1 and VOZ2

To gain insight into possible functions of BNP we carried out a yeast-two-hybrid screen to identify potential BNP protein interactors. Two closely related NAC family transcription factors, VOZ1 and VOZ2, were identified among interactors (all interacting clones are presented in Table S1). Interestingly, VOZ1 and VOZ2 were reported to bind a pollen-specific promoter element (Mitsuda et al., 2004). BNP was also retrieved in the Y2H screening as a self-interactor (Table S1). To confirm BNP-VOZ1, BNP-VOZ2 and BNP-BNP interactions, we performed Bimolecular Fluorescence complementation (BiFC) assays.

We made constructs to express BNP and each of the candidate proteins fused either to the N- or the C-terminal of yellow fluorescent protein (YFP) fragments to generate BNP-N-YFP and VOZ1-C-YFP, VOZ2-C-YFP or BNP-C-YFP fusion constructs. The fusion constructs under control of the *CaMV 35S* promoter were used to transiently transform *N. benthamiana* leaves and BiFC signals were analyzed using confocal microscopy (Fig. 4). YFP signal was detected with VOZ1, VOZ2 and with BNP indicating the interaction of the pairs BNP-VOZ1, BNP-VOZ2 and BNP-BNP (Fig. 4). Control *N. benthamiana* leaves transfected either with BNP-N-YFP, VOZ1-C-YFP, VOZ2-C-YFP or BNP-C-YFP did not show any BiFC signal (Fig. 4). An additional control experiment using VLG (as BNP closest homolog, sharing 72.0% identity) did not show an interaction with VOZ1, VOZ2 and BNP indicating the specificity of the observed interactions (Fig. S5).

**Figure 4.**
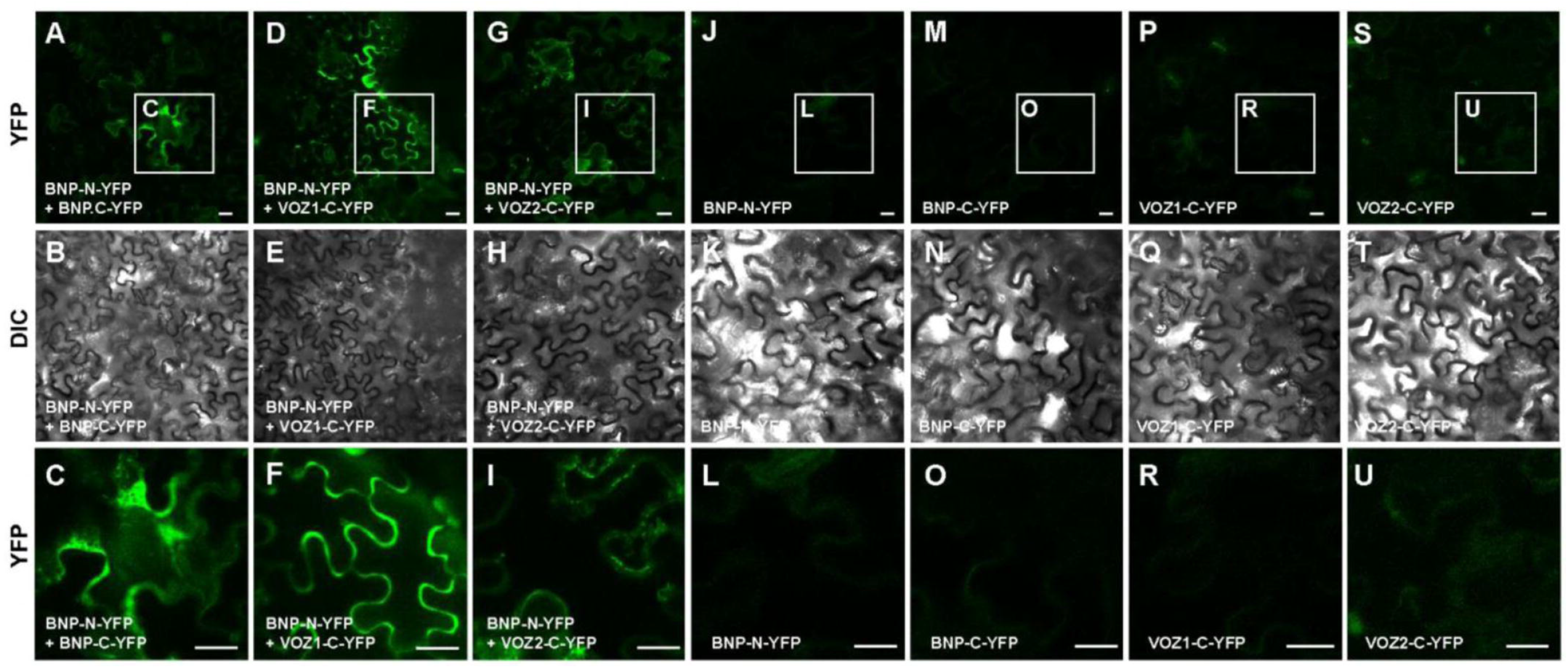
BNP interacts with itself, VOZ1 and VOZ2. Bimolecular fluorescence complementation (BiFC) analysis of the interaction of BNP with itself, VOZ1 and VOZ2 in *N. benthamiana* leaves. Representative confocal microscopy images showing that BNP interacts with (A–C) itself, (D-F) VOZ1 and (G-I) VOZ2. Reconstituted YFP fluorescence (YFP) or DIC images of *N. benthamiana* leaves are shown in epidermal cells co-infiltrated with *Agrobacterium tumefaciens* harboring the indicated constructs. Control *N. benthamiana* leaves transfected with (J-L) BNP-N-YFP, (M-O) BNP-C-YFP, (P-R) VOZ1-C-YFP, (S-U) VOZ2-C-YFP do not show any BiFC signal. The last row (C, F, I, L, O, R, U) correspond to the magnification of regions of interest (insets) indicated in the first row. Bars = 20 μm.

### BNP mainly localizes to Rha1 positive vesicles and co-localizes with VOZ1 and VOZ2

We used the lipophilic dye FM4-64 to gain insight on the subcellular localization of BNP. After internalization, this dye is sequentially distributed to different organelle membranes, following the endocytic pathway (Bolte et al., 2004). We followed FM4-64 in roots of plants expressing the fusion protein BNP-GFP at different time points. We observed no co-localization of internalized FM4-64 and BNP-GFP at early time points of incubation (10-30 min) discarding localization of BNP-GFP to early endosomes or to components of the *trans*-Golgi Network (Fig. S6 A-E). However, high levels of co-localization were observed at about 120 min of incubation, supporting a localization in late endosomes (Fig. S6 F-J). In agreement with that, BNP-GFP did not co-localize with Brefeldin bodies caused by the addition of the endosomal recycling inhibitor Brefeldin A to FM4-64 treated roots (Fig. S6 K-O). In addition, treatment with the PI3K inhibitor LY294002, which was reported to cause swelling and association of late endosomes (Takáč et al, 2013) caused a significant increase in the area of BNP compartments (Fig. S7).

To assess BNP localization, we used *N. benthamiana* to transiently express BNP-GFP together with a late endosome/prevacuolar compartments/multivesicular bodies marker, the Rab5 GTPase Rha1 fused to RFP (Foresti et al., 2010) or with a Golgi apparatus marker, the rat sialyltransferase St fused to RFP (Wee et al., 1998). Representative images shown in Fig. 5 and Pearson and Spearman correlation (PSC) values indicated that BNP-GFP co-localized with Rha1-RFP (Fig. 5A-E) and did not co-localize with St-RFP (Fig. 5F-J), suggesting that BNP is mainly confined to late endosome/prevacuolar compartments/multivesicular bodies associated vesicles.

**Figure 5.**
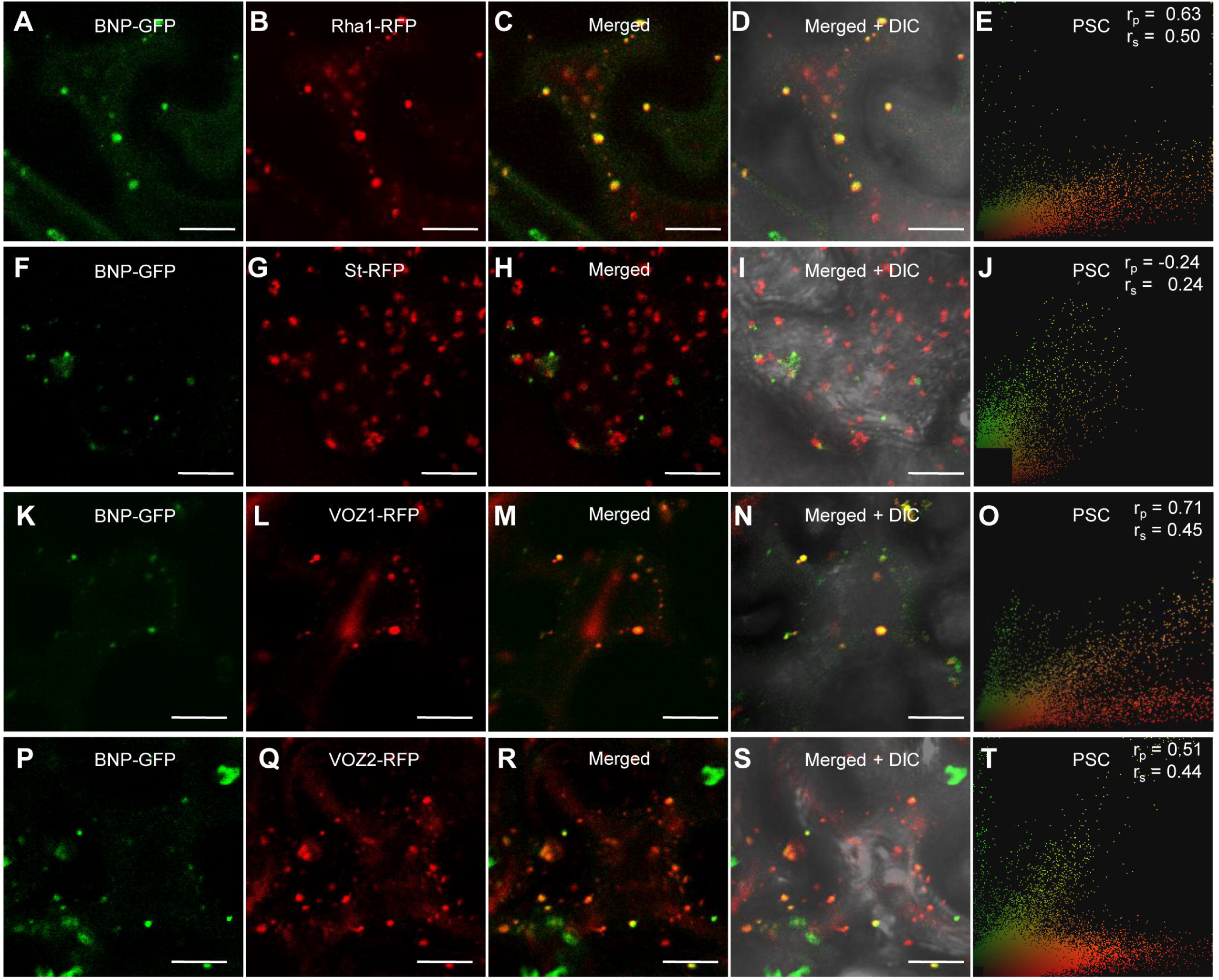
BNP co-localizes with VOZ1 and VOZ2 to Rha1 positive vesicles. (A, F, K and P) Localization of BNP-GFP transiently expressed in *N. benthamiana* epidermal cells. (B) Fluorescent marker for late endosomes/prevacuolar compartments/ multivesicular bodies, RFP-Rha1. (C) Merged image of BNP-GFP and RFP-Rha1. (D) Shows an overlay with the corresponding bright-field image. (E, J, O and T) Pearson and Spearman correlation (PSC) test calculated using a plugin for ImageJ; r values indicate the level of co-localization ranging from +1 for perfect co-localization to -1 for negative correlation. (G) Fluorescent marker for Golgi apparatus, St-RFP. (H) Merged image of BNP-GFP and St-RFP. (I) Shows an overlay with the corresponding bright-field image. (L) Localization of RFP-VOZ1. (M) Merged image of BNP-GFP and RFP-VOZ1. (N) Shows the corresponding bright field image of the cells showed in (M). (Q) Localization of RFP-VOZ2. (R) Merged image of BNP-GFP and RFP-VOZ2. (S) Bright field image of the cells showed in (R). Bars = 20 μm.

To further confirm that BNP and VOZ proteins are indeed localized in the same subcellular compartment, we performed a co-localization study. *N. benthamiana* plants were transiently co-transfected with constructions to express BNP-GFP and either VOZ1-RFP or VOZ2-RFP. In these transfected cells BNP-GFP and VOZ1-RFP or VOZ2-RFP can be found separately, however, a significant shared pixel population indicates partial co-localization, confirmed by moderate correlation scores for Pearson and Spearman tests (Fig. 5K-T).

### BNP is required for nuclear localization of VOZ1 in pollen

We next analyzed VOZ1 localization in pollen at anthesis from *BNP/BNP* and hemizygous *bnp-1/BNP* plants (Fig. 6). We observed that in mature trinucleated pollen VOZ1 localizes to the cytoplasm and to the vegetative nucleus, however, in mature but binucleated pollen from *bnp-1/BNP* plants, VOZ1 is localized only in the cytoplasm (Fig. 6). This result suggested that the absence of a functional BNP could be affecting the localization of VOZ to the nucleus

**Figure 6.**
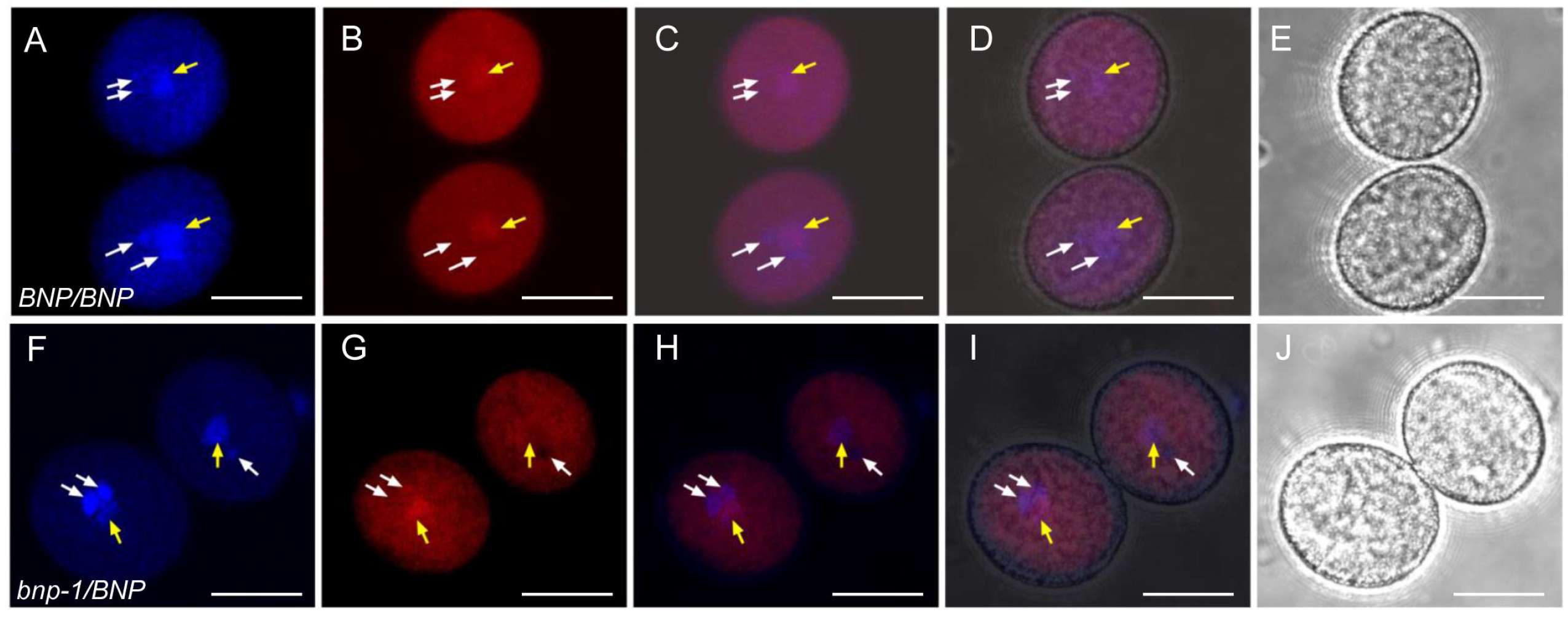
VOZ1 localization in mature pollen from WT and *bnp-1/BNP* plants. Mature pollen from (A- E) BNP/BNP, VOZ1-RFP and (F – J) bnp-1/BNP, VOZ1-RFP plants was analized. Representative z-stack images of confocal micrographs displaying (A, F) DAPI nuclear staining, (B, G) VOZ1-RFP fluorescence and (E, J) bright-field images. (C, H) Merged DAPI and VOZ1-RFP fluorescence images and (D, I) merged DAPI, VOZ1-RFP fluorescence and bright-field images Yellow arrows indicate the vegetative nucleus and white arrows indicate the generative nucleus. Bars = 10 μm.

To test this, we transformed *BNP/BNP* and hemizygous *bnp-1/BNP* plants, both in the *qrt1* background, with *ProUb10:VOZ1-RFP* and analyzed VOZ1-RFP intracellular localization during pollen development (Fig. 7). The introduction of the mutation in the *qrt1* background allows to compare and analyze any difference in VOZ1 localization that might arise from an impairment in BNP, as pollen tetrads of the *bnp-1/BNP* genotype are constituted by two mutant (binucleate) and two WT (trinucleate) pollen cells.

**Figure 7.**
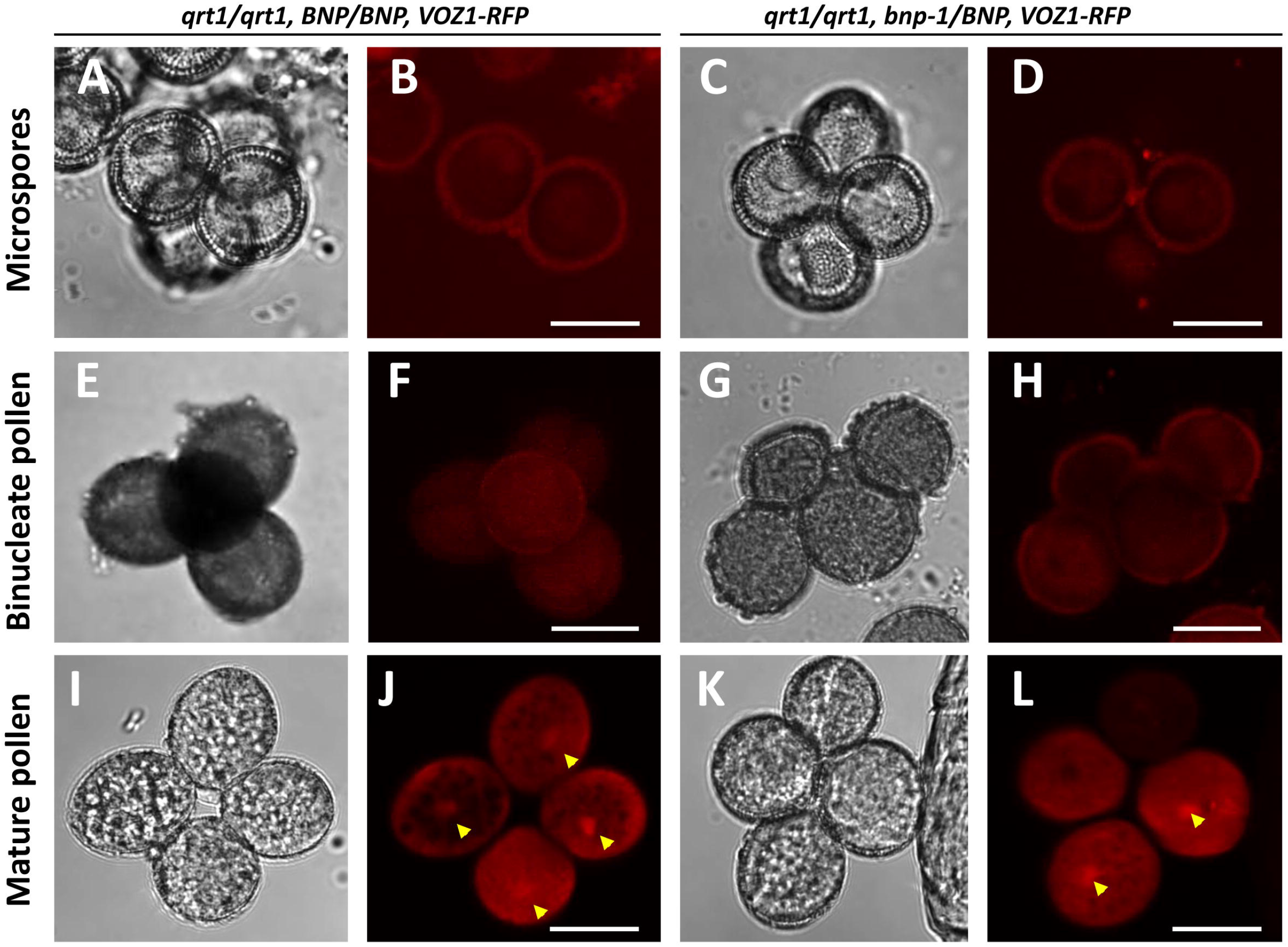
Nuclear localization of VOZ1 is affected in *bnp* mutants. Pollen tetrads from *qrt1/qrt1, BNP/BNP, VOZ1-RFP* (A, B, E, F, I, J) and *qrt1/qrt1, bnp-1/BNP, VOZ1-RFP* (C, D, G, H, K, L) plants were analyzed at microspore (A-D), binucleate (E-H) and mature tricellular (I-L) stages. Representative z-stack images of confocal micrographs displaying VOZ1-RFP fluorescence (B, D, F, H, J, L) and bright field images (A, C, E, G, I, K) are shown. Arrowheads indicate nuclear localized VOZ1. Bars = 10 μm.

At earlier developmental stages (microspore and bicellular pollen) VOZ1 was detected at the cytoplasm with very low signal intensities, showing no differences between *qrt1/qrt1, BNP/BNP, VOZ1-RFP* and *qrt1/qrt1, bnp-1/BNP, VOZ1-RFP* genotypes (Fig. 7A-H). In mature WT pollen, however, VOZ1-RFP was detected at higher levels and its localization appeared distributed between the cytoplasm and the nuclei (Fig. 7I-L). Most of the pollen tetrads (91.5%) from *qrt1/qrt1, BNP/BNP, VOZ1-RFP* plants presented VOZ1 nuclear localization in the four cells of the tetrad (Table 4). However, most of the tetrads from the *qrt1/qrt1, bnp-1/BNP,* VOZ1-RFP genotype presented VOZ1 nuclear localization only in one or two pollen cells (85.2%, Table 4).

**Table 4.** VOZ1 localization in mature pollen in *bnp* mutants expressing VOZ1. Mature pollen tetrads from *qrt1/qrt1, BNP/BNP, VOZ1-RFP* and *qrt1/qrt1, bnp-1/BNP, VOZ1-RFP* plants.

| Genotype | Observed VOZ1 localization |  |  |  |  |
| --- | --- | --- | --- | --- | --- |
|  | Four nuclear | Three nuclear, one cytoplasmic | Two nuclear, two cytoplasmic | One nuclear, three cytoplasmic | Four cytoplasmic |
| *qrt1/qrt1*, *BNP/BNP*, *VOZ1-RFP* | 91.5 % (65/71) | 7.0 % (5/71) | 1.4 % (1/71) | 0 % (0/71) | 0 % (0/71) |
| *qrt1/qrt1*, *bnp-1/BNP*, *VOZ1-RFP* | 1.3 % (1/76) | 11.8 % (9/76) | 57.8% (44/76) | 27.6% (21/76) | 1.3 % (1/76) |

In parenthesis, number of individuals observed over the total analyzed.

Nuclear composition of mature pollen from plants expressing VOZ1-RFP *(qrt1/qrt1, BNP/BNP, VOZ1-RFP* and *qrt1/qrt1, bnp-1/BNP, VOZ1-RFP)* shown in Table 5 presented comparable values to the results showed in Fig. 2A for *qrt1/qrt1, BNP/BNP* and *qrt1/qrt1, bnp-1/BNP* respectively.

**Table 5.** Nuclear composition of mature pollen in *bnp* mutants expressing VOZ1. Mature pollen tetrads from *qrt1/qrt1, BNP/BNP, VOZ1-RFP* and *qrt1/qrt1, bnp-1/BNP, VOZ1-RFP* plants.

| Genotype | Observed nuclear phenotype |  |  |  |
| --- | --- | --- | --- | --- |
|  | Four trinucleate | Three trinucleate, one binucleate | Two trinucleate, two binucleate | Two binucleate, one tetranucleate |
| *qrt1/qrt1*, *BNP/BNP*, *VOZ1-RFP* | 97.3 % (71/73) | 2.7 % (2/73) | 0 % (0/73) | 0 % (0/73) |
| *qrt1/qrt1*, *bnp-1/BNP*, *VOZ1-RFP* | 3.6 % (4/110) | 18.2 % (20/110) | 77.3 % (85/110) | 0.9 % (1/110) |

In parenthesis, number of individuals observed over the total analyzed.

Next, we analyzed if the lack of a functional BNP could also affect the levels of VOZ1 in pollen. We quantified VOZ1-RFP by measuring red fluorescence intensity in mature pollen cells of *qrt1/qrt1, BNP/BNP,* VOZ1-RFP and *qrt1/qrt1, bnp-2/BNP,* VOZ1-RFP plants (Fig. S8). Interestingly, lower red fluorescence levels were recorded in *qrt1/qrt1, bnp-2/BNP,* VOZ1-RFP plants suggesting that BNP could have a role in the promotion of VOZ1 stabilization. Altogether, these results strongly suggest that BNP is involved in VOZ1 localization, as its nuclear localization relies on the presence of a functional BNP.

We then analyzed BNP localization during pollen development using WT plants harboring stable *BNP-GFP* and *VOZ1-RFP* constructs. We observed that both, VOZ1 and BNP, localized in the cytoplasm at early developmental stages. Contrasting VOZ1, no nuclear localization was recorded for BNP (Fig. S9). Partial co-localization was detected in areas in the cytosol were BNP and VOZ1 might interact at microspore and bicellular stages. This was not observed in mature pollen, where VOZ1 show both nuclear and cytoplasmic localization. As BNP remained in the cytosol at every developmental stage, forming speckles during binucleate and mature pollen stages, any role related to VOZ proteins translocation/function would be performed outside the nucleus and before pollen maturation.

## 3. Discussion

In this study we identified *BNP,* a gene coding for a DC1 domain protein that is required for pollen development and germination. Analysis of hemizygous *bnp/BNP* plants showed that transmission through the male gametophyte is severely impaired; about half of pollen grains from *bnp/BNP* plants resulted arrested by the binucleate stage and their germination was significantly reduced.

By means of Y2H and BiFC experiments, we identified and confirmed VOZ1 and VOZ2 as BNP interacting proteins. VOZ1 and VOZ2 are transcription factors that belong to the NAC transcription factor family, and were reported as involved in the control of flowering time (Celesnik et al., 2013; Yasui and Kohchi, 2014), either through the binding of PhyB (Yasui et al., 2012), Flowering Locus C (Mimida et al., 2011) or Constans (Koguchi et al., 2017; Kumar et al., 2018). VOZ proteins are also involved in biotic and abiotic stress responses (Neill et al., 2002; Nakai et al., 2013; Song et al., 2018; Schwarzenbacher et al., 2020). VOZ1 and VOZ2 were proved to be functionally redundant, as single mutants for each gene showed normal phenotypes, but *voz1voz2* double mutants displayed phenotypic defects (Yasui et al., 2012; Celesnik et al., 2013; Nakai et al., 2013; Kumar et al., 2018). Interestingly, *voz1voz2* plants presented the same pollen developmental defect observed in *bnp/BNP* plants, as a fraction of their pollen grains contained only two nuclei –the vegetative nucleus and a single reproductive nucleus (Celesnik et al., 2013). This fact supports a physiological role for the interaction between BNP and VOZ proteins during pollen development.

BNP was localized mainly in Rha1 positive vesicles, which are defined as late endosomes/ prevacuolar compartments/ multivesicular bodies. VOZ proteins are functional in the nucleus, but they were initially identified in cytoplasmic speckles and a cytoplasmic-nuclear translocation was proposed to modulate VOZ activity as transcriptional regulators (Yasui et al., 2012; Koguchi et al., 2017). VOZ1 and VOZ2 do not contain neither a transmembrane region nor localization signals such as NLS or NES (Jensen et al., 2010). This suggests that VOZ1 and VOZ2 nuclear/cytoplasmic localization must rely on another mechanism. Endocytic proteins have been previously shown to participate in the spatiotemporal regulation of transcription factors (Palfy et al., 2012; Wu et al., 2012; Serrano et al., 2016). They may facilitate the assembly of multi-molecular complexes through protein–protein interaction domains and act as regulators (Pyrzynska et al., 2009). As can be concluded from the analysis showed in Fig. 6, 7 and Table 3 a similar situation could be taking place between the pairs BNP-VOZ1, where VOZ1 is required to interact with BNP to reach the nuclear import machinery.

The involvement of the endomembrane system in nuclear translocation has been previously reported, as several membrane receptors undergo endocytosis and are transported to the nucleus via endosomal trafficking (Mukherjee et al., 2006, Wang et al., 2010; Wang et al., 2020). However the mechanisms of the process are still under investigation and several mechanisms are proposed to allow the cargo reach the nuclear import machinery, including retrograde movement of vesicles through the Golgi and ER, the release of the cargo in the cytoplasm or the exposing of a NLS that is recognized by importin-β (Ek-Ramos et al., 2014).

Implications of VOZ1/2 nuclear translocation may be related to VOZ1/2 role as a transcriptional regulator, with such regulation occurring in the nucleus (Yasui et al., 2012). Interestingly, VOZ proteins nuclear localization was rarely detected, and the addition of NLS signals was usually required to place it the nucleus (Yasui et al., 2012; Schwarzenbacher et al., 2020) However it was reported that degradation of VOZ1/2 takes also place in the nucleus, facilitated by the E3 ubiquitin ligase BRUTUS (Selote et al., 2018). Thus, BNP role could be involved in the function and/or in the degradation of VOZ1/2.

Our results also indicated that BNP could be essential not only during pollen development but also at additional developmental stages, such as germination and early stages of sporophyte development. In agreement with this, *BNP* expression was detected very early during seedling development, in particular in vascular tissues (Fig. 3). The expression pattern of BNP also suggests that it could play roles in other sporophytic tissues, either through the interaction with VOZ TFs, that are also ubiquitously expressed through the plant (Yasui et al., 2012), or through the interaction with other proteins. Potential BNP interacting proteins identified by Y2H experiments (Table S1) provide a promising list of candidates that might be involved with BNP function in other plant tissues.

Only one additional DC1 domain protein, VLG, has been characterized so far and was likewise proved to be required for pollen development. However, *VLG* deficiency caused microgametogenesis arrest at PMI, an earlier stage, yielding unviable uninucleate pollen grains (D’Ippólito et al., 2017). On the other hand, VLG was also essential for female gametogenesis (D’Ippólito et al., 2017).

Notably, the Arabidopsis genome encodes for over 140 uncharacterized proteins harboring DC1 domains. Multiple sequence alignments of DC1 domain proteins from different species showed amino acid conservation restricted to the residues that define the DC1 signature motif, but divergence in the amino acids corresponding to the folded domain -which might define binding capabilities (Rahman and Das, 2015)- suggesting a diversity of interactors. Thus, DC1 domain proteins emerge as a novel family of highly specialized modular scaffold proteins, capable to promote the recruiting of binding targets and affect their localization and functionality.

Altogether, our results support a role for BNP acting as a scaffold protein recruiting VOZ1 and VOZ2 to the endomembrane system. This recruitment appears to facilitate VOZ1 -and likely VOZ2-translocation to the nucleus, promoting their activity as transcriptional regulators during pollen development. In addition, a role of BNP in the regulation of the levels VOZ proteins could not be discarded, as lower levels of these proteins were present in mature pollen lacking a functional BNP.

## 4. Materials and Methods

### Plant materials and growth conditions

Arabidopsis (Columbia [Col-0] ecotype) lines SALK_114889 *(bnp-1,* with T-DNA insertion located in chr2 TAIR10 18,323,075) and GK-008E01 *(bnp-2,* with T-DNA insertion located in chr2 TAIR10 18,323,371) were obtained from ABRC (Ohio State University, Ohio, USA) (Alonso et al., 2003). Both lines were backcrossed twice to the WT Columbia prior to their use. The presence of a single insertion in *At2g44370* and all genotypes was confirmed by PCR-based genotyping and segregation analysis. When indicated seeds were sterilized in 20% (v/v) sodium hypochlorite, washed with sterile water and plated on MS plates with 50 μg/mL kanamycin or 7.5 μg/mL sulfadiazine, stratified at 4°C for 48 h and placed at 23°C, 16 h light, 8 h dark. Resistant seedlings were then transferred onto soil and grown under the conditions described above.

### Molecular characterization of insertional lines

For *bnp-1,* the left border–genomic sequence junction was determined by PCR in plants showing kanamycin resistance using the T-DNA-specific primer LBb1 combined with the genomic sequence specific primers bnp-1 RP and bnp-1 LP (Table S2). For *bnp-2,* the left border–genomic sequence junction was determined by PCR in plants showing sulfadiazine resistance using the T-DNA–specific primer GKGT8474 combined with the genomic sequence specific primers bnp-2 RP and bnp-2 LP (Table S2).

### Segregation Analysis

For self-fertilization analysis, the progeny of self-pollinated hemizygous plants was germinated on selective growth medium containing 50 μg/mL kanamycin for *bnp-1* or 7.5 μ/mL sulfadiazine for *bnp-2,* and the ratio of resistant to sensitive plants was scored. Reciprocal crosses were performed as described previously (Pagnussat et al., 2005).

### Generation of vectors for plant transformation

Genomic DNA was extracted from rosette leaves or whole seedlings as described (León et al., 2007). To generate *ProBNP:BNP-GFP* translational fusion the region including *BNP* ORF and putative promoter (672 upstream the ATG codon) was amplified by PCR using the primer combination listed in Table S2. The amplicon was cloned using pENTR™ Directional TOPO^®^ Cloning Kit (Invitrogen) using Gateway^®^ technology and the sequence was verified. The resulting plasmid *pENTR-ProBNP-BNP* was subjected to the LR reaction using the destination vector pMDC107 (Curtis and Grossniklaus, 2003). The *ProBNP:GUS* construct was generated amplifying the *BNP* putative promoter region (672 upstream the ATG codon) by PCR using the primer combination listed in Table S2. The amplicon was cloned into pENTR™/TOPO (Invitrogen) and its sequence was verified. The resulting plasmid *pENTR-ProBNP* was subjected to the LR reaction using the destination vector pMDC162 (Curtis and Grossniklaus, 2003). For *Pro35S:BNP-GFP* construct, *BNP* ORF was PCR amplified using as template clone DKLAT2G44370 obtained from ABRC (Ohio State University, Ohio, USA) using the primer combination listed in Table S2. The amplicon was cloned into pENTR™/TOPO (Invitrogen) and its sequence was verified. The resulting plasmid *pENTR-BNP* was subjected to the LR reaction using the destination vector pMDC83 (Curtis and Grossniklaus, 2003). For *ProUb10:VOZ1-RFP* and *ProUb10:VOZ2-RFP* constructions, respective ORFs were PCR amplified using primers listed in Table S2, cloned in pENTR™/TOPO (Invitrogen) and sequence-verified. The resulting plasmids *pENTR-VOZ1* and *pENTR-VOZ2* were subjected to the LR reaction using the destination vector pUBN-RFP-DEST (Grefen et al., 2010).

### Obtaining *bnp/BNP* plants in the *qrt1* background

To obtain *bnp* mutants in *qrt1* background, *bnp-1/BNP* was crossed with *qrt1* homozygous plants. Kanamycin resistant plants from the progeny were selected and allowed to self-fertilize. Homozygous *qrt1* plants harboring *bnp* mutations were selected for further analysis (Johnson-Brousseau and McCormick, 2004).

In a similar way, *qrt1/qrt1-bnp-2/BNP-VOZ1-RFP* plants were obtained crossing *bnp-2/BNP* plants expressing VOZ1-RFP with *qrt1* homozygous plants.

### Transformation of *Agrobacterium tumefaciens* and Arabidopsis

Vectors were introduced into Agrobacterium strain GV3101 by electroporation. Arabidopsis WT plants were transformed using the floral dip method (Clough and Bent, 1998). Transformant plants were selected based on their ability to survive in MS medium with 15 mg. L^-1^ hygromycin. Resistant (green seedlings with true leaves) were then transferred to soil and grown under the conditions described above.

### Morphological and histological analyses

Pollen grain viability was assessed using Alexander’s staining (Alexander, 1969) and fluorescein diacetate (FDA) staining (Heslop-Harrison and Heslop-Harrison, 1970). For pollen development analysis, successive buds in the same inflorescence were collected (Lalanne and Twell, 2002), fixed in ethanol:acetic acid 3:1 until discoloration and stored at room temperature. Anthers were washed, dissected and mounted on microscopy slides with DAPI staining solution. For pollen tube germination, pollen grains were incubated overnight in a dark moisture chamber at 22°C in 0.01% boric acid, 5mM CaCl_2_, 5mM KCl, 1 mM MgSO_4_, 10% sucrose pH 7.5 and 1.5% agarose, as described previously (Boavida and McCormick, 2007). For GUS staining, tissues from *ProBNP:GUS* carrying plants were collected and incubated in GUS staining solution as already described (Pagnussat et al., 2007). Gametophytes were observed on a Zeiss Axio Imager A2 microscope using DIC optics. Seedlings were observed on a Nikon SMZ800 Stereomicroscope and images were captured with a Canon DS126431 camera.

### Yeast two-hybrid screen

The Y2H screening was performed by Hybrigenics Services S.A.S. (Paris, France). A DNA fragment encoding Arabidopsis At2g44370 (amino acids 1-250) was PCR-amplified and cloned into pB66 as C-terminal fusion to Gal4 DNA-binding domain (Gal4-At2g44370) and used as a bait to screen Universal Arabidopsis Normalized library containing 3.2 million of independent clones in pGADT7-RecAB vector. pB66 derive from the original pAS2ΔΔ (Fromont-Racine et al., 1997). For the Gal4 bait construct, 35 million clones were screened using a mating approach with YHGX13 (Y187 ade2-101::loxP-kanMX-loxP, matα) and L40ΔGal4 (mata) yeast strains as previously described (Fromont-Racine et al., 1997). 158 His+ colonies were selected on a medium lacking tryptophan, leucine and histidine. The prey fragments of the positive clones were amplified by PCR and sequenced at their 5’ and 3’ junctions. The resulting sequences were used to identify the corresponding interacting proteins in the GenBank database (NCBI) using a fully automated procedure, and in total, 23 different clones were identified (Table S1). A confidence score (PBS, for Predicted Biological Score) was attributed to each interaction as previously described (Formstecher et al., 2005).

### Co-localization experiments in *Nicotians benthamiana*

*A. tumefaciens* strain GV3101 carrying *Pro35S:BNP-GFP* was co-infiltrated with *A. tumefaciens* cells carrying either the late endosome marker *RFP-Rha1* or the Golgi apparatus marker *St-RFP,* in leaf epidermal cells of *N. benthamiana* plants. Infiltrated leaves were analyzed 48-72 h after infiltration on a confocal microscope (Nikon Eclipse C1 Plus) using EZ-C1 3.80 imaging software and Ti-Control. Pearson and Spearman correlation (PSC) plugin for ImageJ was used to calculated correlation coefficients from a minimum of 400 individual regions of interest corresponding with punctate structures were manually selected from at least 10 independent cell images (French et al., 2008).

### Co-localization experiments in Arabidopsis

Arabidopsis root cells expressing a *35S:BNP-GFP* construction were used to analyze the co-localization between BNP and the endocytic tracer FM4-64. Roots were incubated for the indicated times with 6 μM FM4-64 solution, washed twice, and then incubated in liquid medium or with the indicated inhibitors. For Brefeldin A treatments, incubation was performed in 50 μM Brefeldin A solution for 15 min. For LY294002 treatments, incubation was performed in 100μM LY294002 solution for 120 min. Confocal microscopy images were obtained at the indicated times with a confocal microscope (Nikon Eclipse C1 Plus) using EZ-C1 3.80 imaging software and Ti-Control. Pearson and Spearman correlation (PSC) plugin for ImageJ was used to calculated correlation coefficients from a minimum of 400 individual regions of interest from at least 10 independent cell images (French et al., 2008).

### Bimolecular fluorescence complementation (BiFC) analysis

The cDNA sequences of At2g44370 (*BNP*), At1g28520 (*VOZ1*), At2g42400 (*VOZ2*) and At2g17740 (*VLG*) were PCR amplified using the primers listed in Table S2. The PCR products were cloned into pENTR™/TOPO (Invitrogen), sequenced and recombined through BP reaction into BiFC destination plasmids pUBN-YN and pUBN-YC (Grefen et al., 2010). The binary plasmids were then transformed into *A. tumefaciens* strain GV3101 by electroporation. Split nYFP- and cYFP-tagged protein pairs and the gene-silencing suppressor p19 were co-expressed in *N. benthamiana* leaves by *A. tumefaciens* -mediated inoculation. Plant leaves were examined 48-72 h post-infiltration with a confocal microscope (Nikon Eclipse C1 Plus) using a Super Fluor 40.0x/1.30/0.22 Oil objective, numerical aperture 1.300. Acquisition was performed using EZ-C1 3.80 imaging software and Ti-Control. All pictures were acquired with the same settings.

### Quantification of VOZ1-RFP in mature pollen

High quality images of mature pollen grains from the genotypes *qrt1/qrt1, BNP/BNP, VOZ1-RFP* and *qrt1/qrt1, bnp-1/BNP, VOZ1-RFP* plants were acquired and red fluorescence was quantified. Red fluorescence background of mature pollen grains from plants not expressing VOZ1-RFP was subtracted. Fluorescence was expressed in arbitrary units relative to the calculated cell area of each pollen grain. Data points were plotted on a violin plot using GraphPad, median (blue line) and quartiles (black lines) are indicated. Box: Mean values ± SD and number of data points (n) is indicated. A significant difference of the means was indicated using unpaired t-test.

### Bioinformatics and phylogenetic analysis

Sequences from the DC1 domain family were identified using the Basic Local Alignment Search Tool (BLAST) from NCBI (National Center for Biotechnology Information) using initially BNP as a reference and retrieved sequences in iterative searches. Protein sequences alignments were performed using MEGA7 (version 7.0.14) (Kumar et al., 2016). Phylogenetic trees were constructed using the neighbor-joining method and the default settings of MEGA7 (version 7.0.14) (Kumar et al., 2016). The evolutionary distances were computed using the Poisson correction method (Zuckerkandl and Pauling, 1965) and are in the units of the number of amino acid substitutions per site. The trees were drawn using FigTree (version 1.4.3) (http://tree.bio.ed.ac.uk/). Graphic display of identities was visualized using Geneious (version 9.1.4) (http://www.geneious.com) based on an identity matrix (Kearse et al., 2012).

## 5. Data availability

All newly generated sequences are available for peer-review and registered in public repositories. Sequence data from this article can be found in the Arabidopsis Genome Initiative database under accession numbers: BNP (At2g44370), VOZ1 (At1g28520), VOZ2 (At2g42400) and VLG (At2g17740).

## 7. Acknowledgements

We would like to thank Michael Blatt and Christopher Grefen (Glasgow University, UK) for providing pUB-Destination vectors, Jurgen Denecke (University of Leeds, UK) for providing the constructs ST-RFP and Rha-RFP, Nicolas Setzes for assistance with statistical analysis and Daniela Villamonte for technical assistance with confocal microscopy. L.A.A., J.F. and F.M. are fellows of CONICET; N.L.A. is a fellow of ANPCyT; D.F.F., G.C.P., C.A.C. and S.D. are researchers from CONICET.

## 8. Author contributions

D.F.F. conceived the original screening and research plans; D.F.F., G.C.P. and C.A.C. supervised the experiments; L.A.A., S.D., J.F., N.L.A and F.M. performed most of the experiments; D.F.F., G.C.P and L.A.A. designed the experiments and analyzed the data; D.F.F. conceived the project and wrote the article with contributions of all the authors; D.F.F. agrees to serve as the author responsible for contact and ensures communication

