## Supplementary Figures 1-8 and Tables 1-2 for "The DC1 domain protein BINUCLEATE POLLEN is required for pollen development in *Arabidopsis thaliana*"

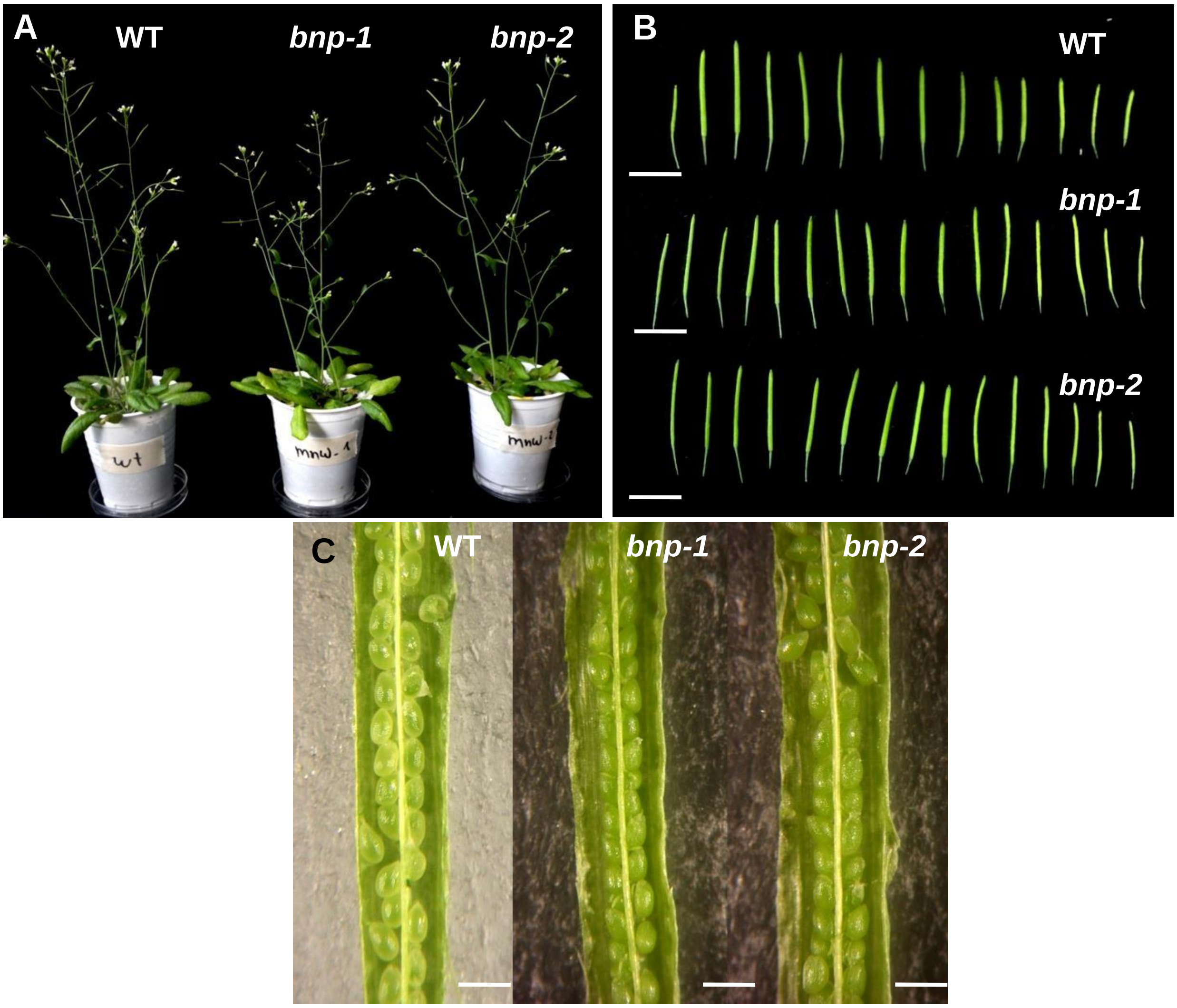


**Figure S1**. ***bnp/BNP* hemizygous plants showed normal sporophytic phenotypes and seed sets.** (A) 5 weeks old WT, *bnp/1-BNP* and *bnp-2/BNP* plants. (B) Siliques of WT, *bnp-1/BNP* and bnp-*2/BNP* plants. (C) Dissected siliques of WT, *bnp-1/BNP* and *bnp-2/BNP* plants at a comparable stage. Bars = 10 mm (B) and 100 μm (C).

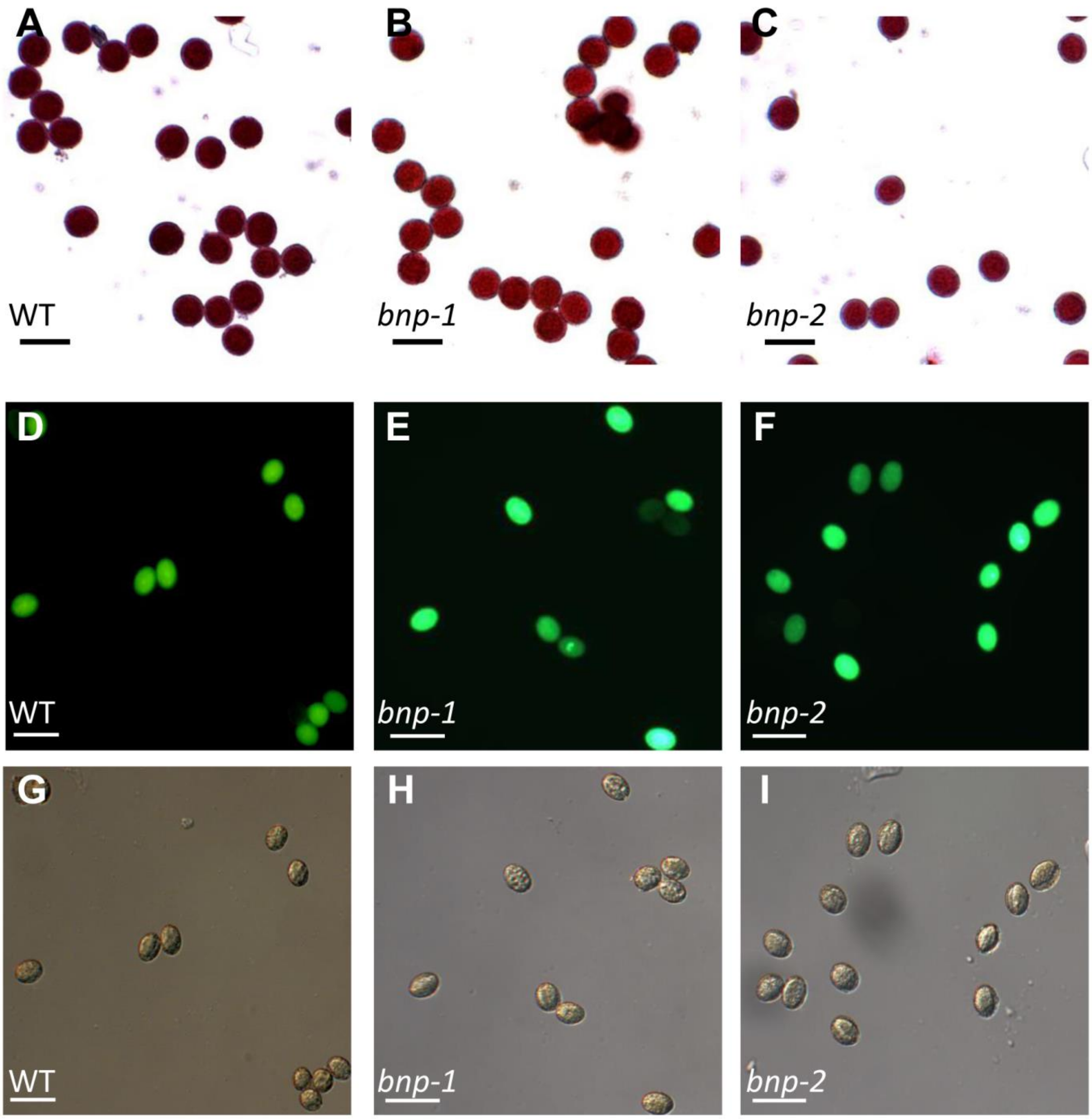


**J**

| Genotype | % Viable pollen (n) |
|---|---|
| WT | 78.3 (428) |
| *bnp-1/BNP* | 77.9 (480) |
| *bnp-2/BNP* | 79.2 (404) |

**Fig. S2**. ***bnp/BNP*** **hemizygous plants showed no differences in the viability of mature pollen.** Pollen viability was analyzed in (A, D, G) WT, (B, E, H) *bnp-1* and (C, F, I) *bnp-2* by means of (A-C) Alexander staining and (D-I) FDA staining. Representative images of (D-F) fluorescence and (G-I) bright-field views are shown. (J) Quantification of pollen viability by means of FDA staining. Bars = 20 µm.

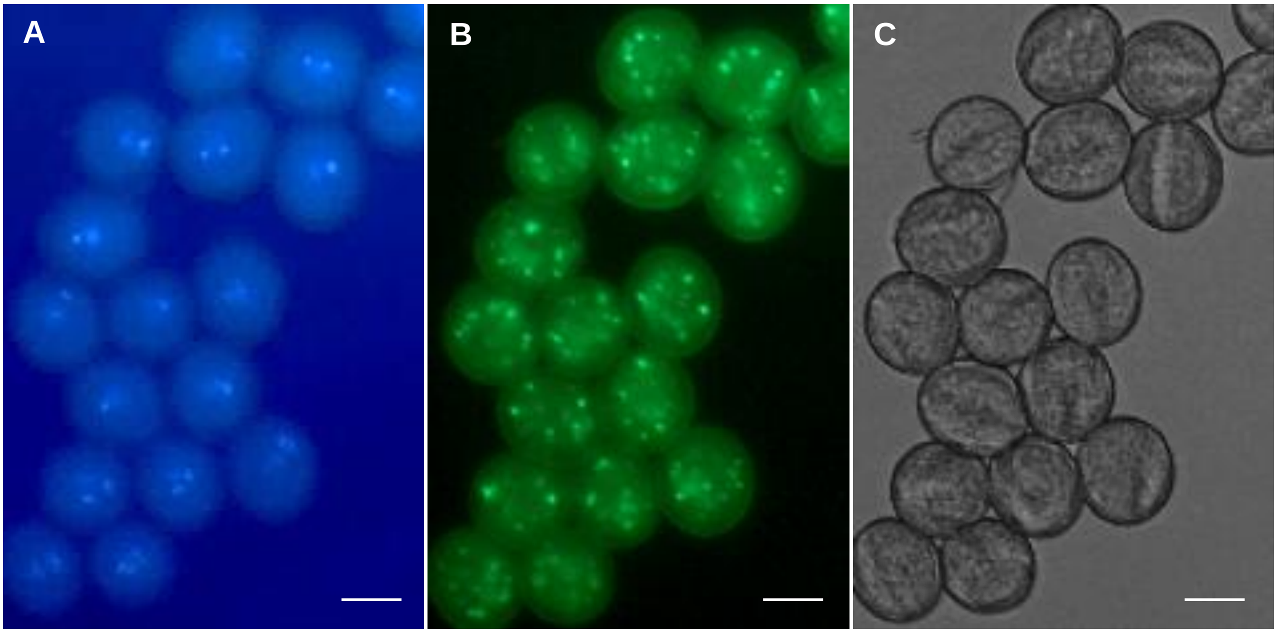


D

| Genotype | % trinucleate pollen (n) |
|---|---|
| WT | 99.2 (772) |
| *bnp-1/BNP, pBNP-BNP-GFP* | 99.3 (860) |

**Figure S3**. **Complementation of *bnp-1/BNP* with the construction *pBNP-BNP-GFP* fully restores pollen binucleate phenotype**. *bnp-1/BNP* plants were transfected with the construction *pBNP-BNP-GFP.* Mature pollen was analyzed by (A) DAPI staining and (B) detection of GFP signal. (C) Bright field image. Pollen grains showed normal nuclear configuration and BNP-GFP fusion protein as detected. (C) Quantification of trinucleate pollen in the complemented line. Bars = 10 µm.

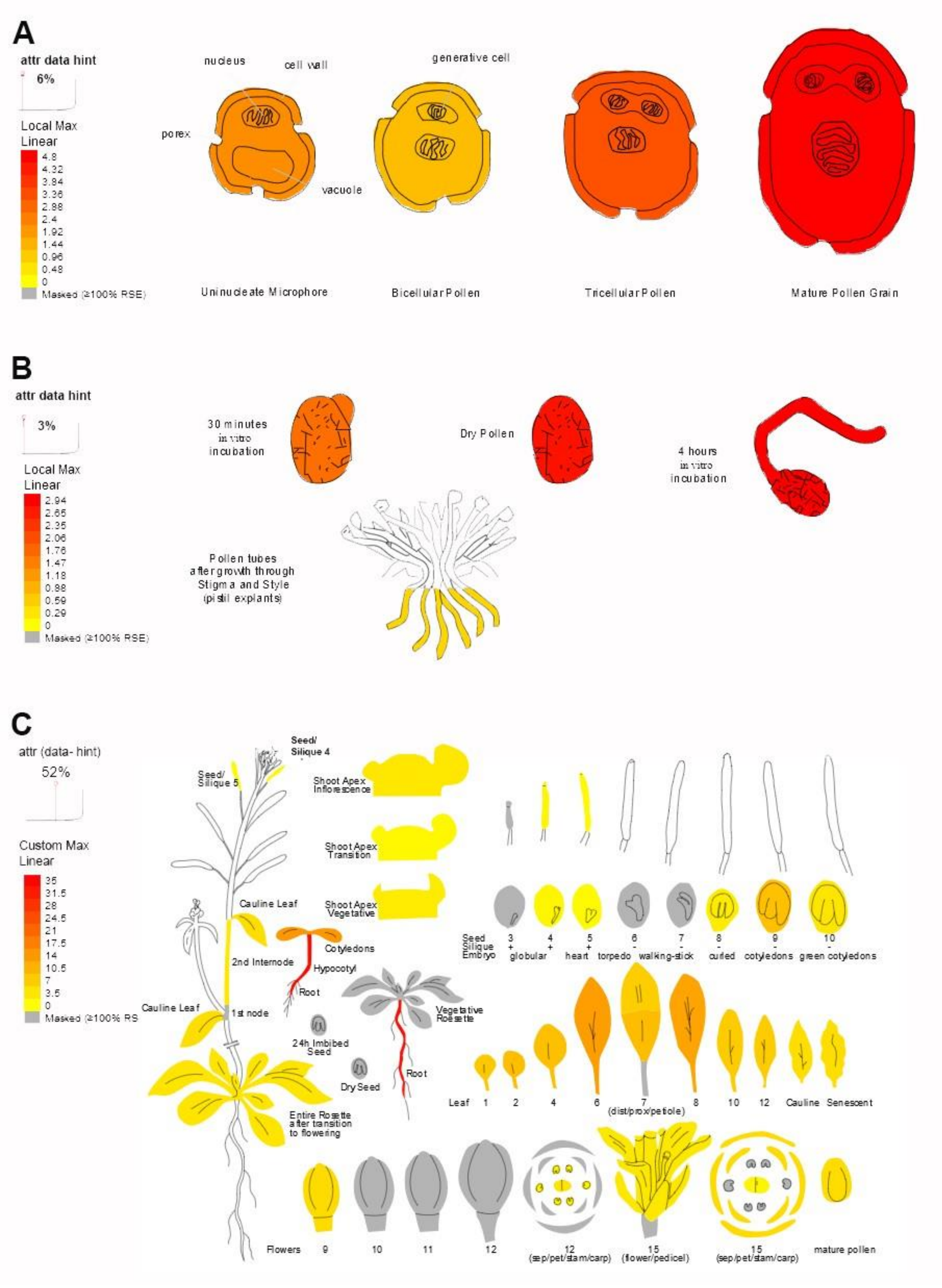


**Figure S4**. **BNP expression in Arabidopsis.** BNP expression during (A) microgametogenesis, (B) pollen germination and (C) in sporophitic tissues. Images generated with the AtGenExpress eFP at bar.utoronto.ca/eplant (Waese et al. 2017). Different scales were used in each figure.

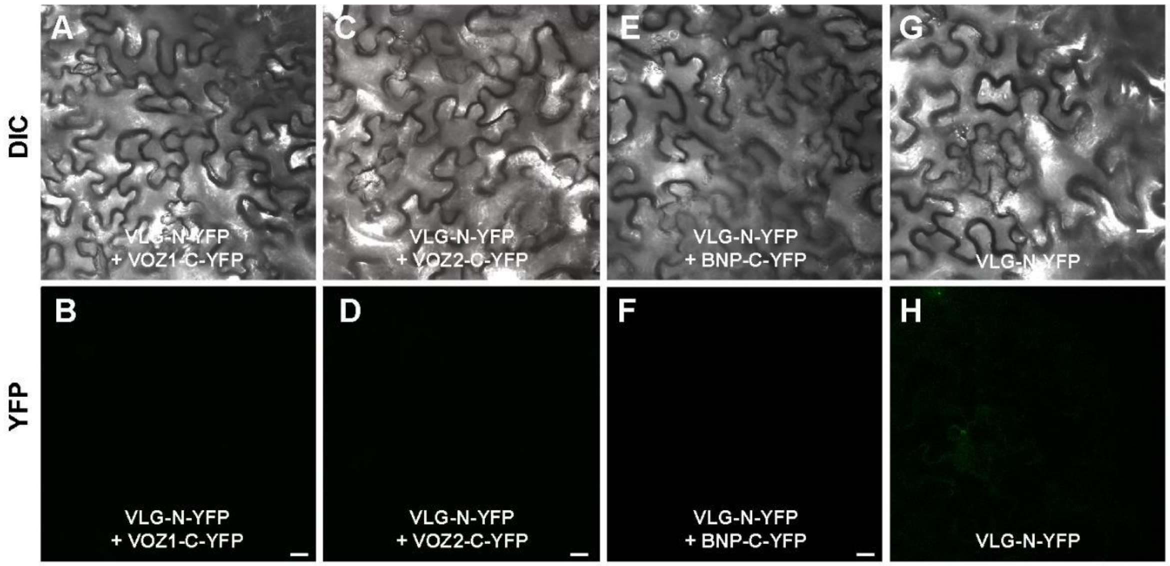


**Figure S5. BNP-related protein VLG does not interacts with VOZ1, VOZ2 and BNP.** Bimolecular fluorescence complementation (BiFC) analysis of the interaction of BNP with VOZ1, VOZ2 and BNP in *Nicotiana benthamiana* leaves. Representative images from confocal microscopy showing that (A-B) VOZ1, (C-D) VOZ2 and (E-F) BNP do not interact with the BNP-related protein VLG. (G, H) Control *N. benthamiana* leaves transfected with VLG-N-YFP do not show any BiFC signal. (A, C, E, G) Reconstituted DIC or (B, D, F, H) YFP fluorescence images of *N. benthamiana* leaves are shown in epidermal cells co-infiltrated with *A. tumefaciens* harboring the indicated constructs. Other negative controls are shown in Fig. 4. Bars = 20 μm.

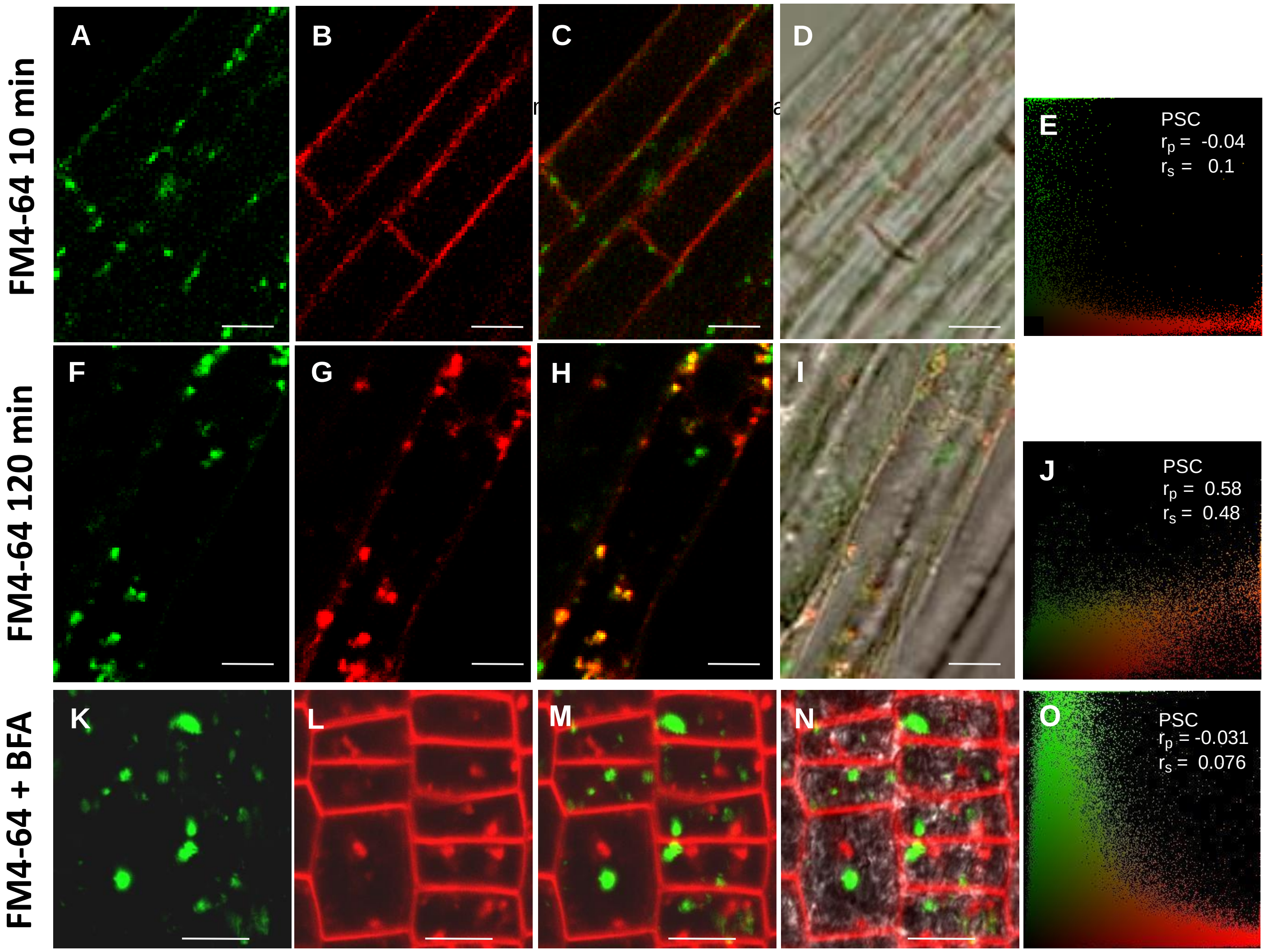


**Figure S6. Analysis of co-localization of BNP and the endocytic tracer FM4-64**. Confocal microscopy images of Arabidopsis root cells expressing a 35S:BNP-GFP construction. Roots were incubated for 5 min with 6 µM FM4-64 solution, washed twice, and incubated in liquid medium (A-E) immediately or (F-J) for 120 min. (K-O) Roots were incubated with 6 µM FM4-64 for 15 min, washed and treated with 50 µM Brefeldin A (BFA) and analyzed after 15 min. (A, F, K) Localization of BNP-GFP, (B, G, L) localization of FM4-64. (C, H, M) Merged green and red fluorescence images, (D, I, N) bright field image with overlayed fluorescence images. (E, J, O) Pearson and Spearman correlation test (PSC) plugin for ImageJ, r values indicate the level of co- localization. Bars = 10 µm.

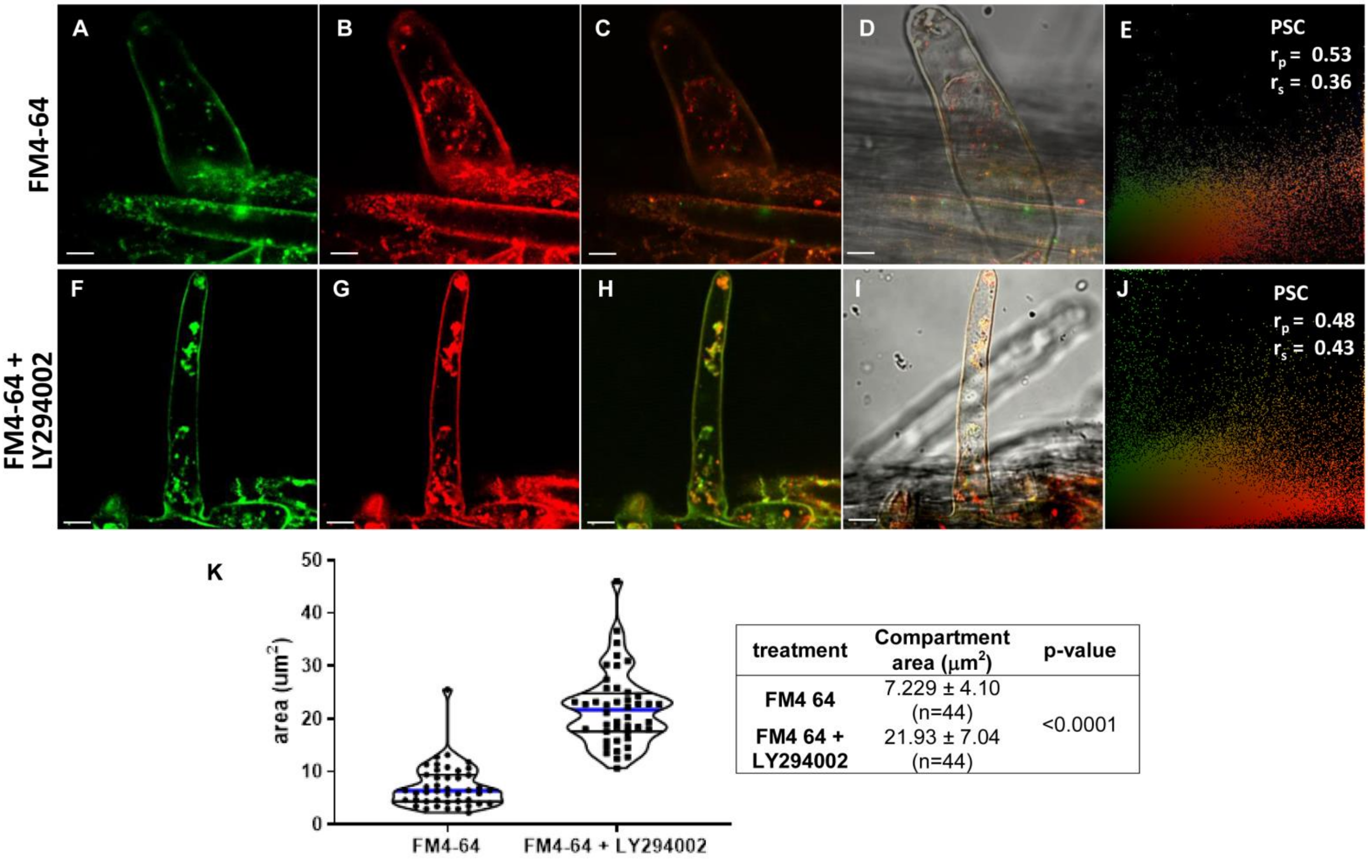


| treatment | Compartment area ($\mu m^2$) | p-value |
|---|---|---|
| FM4 64 | 7.229 ± 4.10 (n=44) | <0.0001 |
| FM4 64 + LY294002 | 21.93 ± 7.04 (n=44) | |

**Figure S7. Analysis of co-localization of BNP and FM4-64 upon LY294002 treatment**. Confocal microscopy images of Arabidopsis root cells expressing a 35S:BNP-GFP construction. Roots were incubated for 5 min with 6 µM FM4-64 solution, washed twice, and incubated (A-E) in liquid medium or (F-J) in 100µM LY294002 for 120 min. (A, F) Localization of BNP-GFP, (B, G) localization of FM4-64. (C, H) Merged green and red fluorescence images, (D, I) bright field image with overlayed fluorescence images. (E, J) Pearson and Spearman correlation test (PSC) plugin for ImageJ between BNP-GFP and FM4-64 signals, r values indicate the level of co- localization. (K) Quantification of BNP compartments in roots treated with FM4-64 or FM4-64 + LY294002. Cell area was calculated and data points were plotted on a violin plot using GraphPad, median (blue line) and quartiles (black lines) are indicated. Box: Mean values ± SD and number of data points (n) is indicated. A significant difference of the means was indicated using unpaired t-test. Bars = 10 µm.

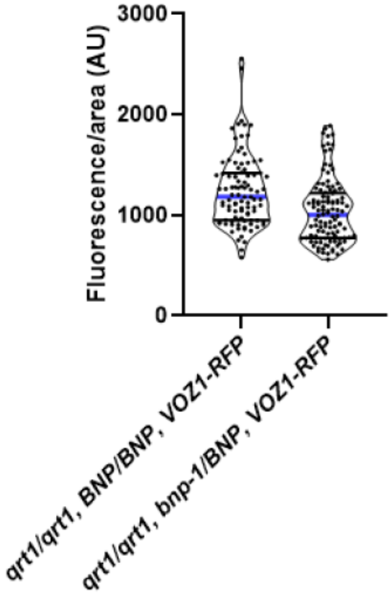


| Genotype | Red fluorescence / cell area (AU) | n | p-value (unpaired t-test) |
|---|---|---|---|
| *qrt1/qrt1, BNP/BNP, VOZ1-RFP* | 1239.75 ± 36.75 | 95 | <0.0001 |
| *qrt1/qrt1, bnp-1/BNP, VOZ1-RFP* | 1042.99 ± 30.41 | 110 | |

**Figure S8. Quantification of VOZ1-RFP red fluorescence in mature pollen of *qrt1/qrt1, BNP/BNP, VOZ1-RFP* and *qrt1/qrt1, bnp-1/BNP, VOZ1-RFP* plants**
High quality images of mature pollen grains from each genotype were acquired and red fluorescence was quantified. Red fluorescence background of mature pollen grains from plants not expressing VOZ1-RFP was subtracted. Fluorescence was expressed in arbitrary units relative to the calculated cell area of each pollen grain. Data points were plotted on a violin plot using GraphPad, median (blue line) and quartiles (black lines) are indicated. Box: Mean values ± SD and number of data points (n) is indicated. A significant difference of the means was indicated using unpaired t-test.

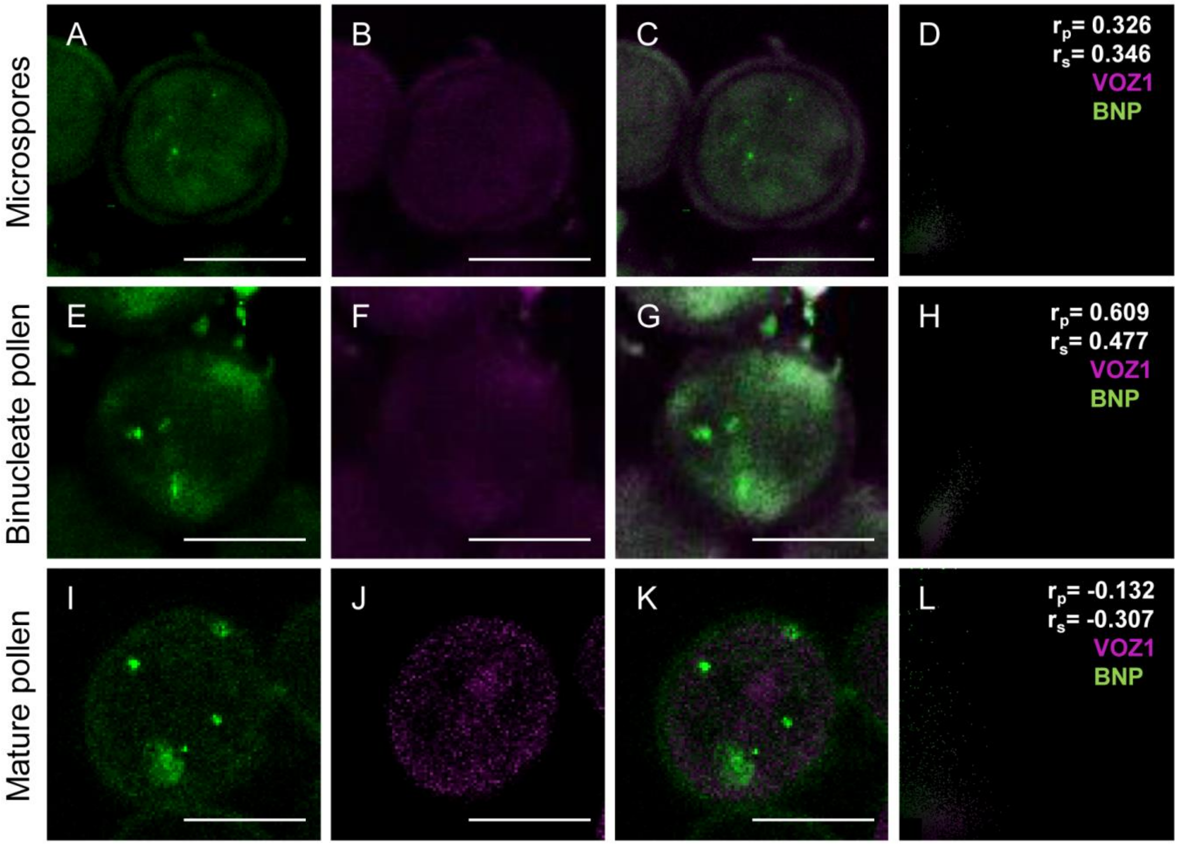


**Figure S9: BNP and VOZ1 localization during pollen development.**
(A-D) Microspore, (E-H) binucleate and (I-L) trinucleate pollen from WT plants harboring stable BNP-GFP and VOZ1-RFP constructs were analyzed. Representative z-stack images of confocal micrographs displaying (A, E, I) BNP- GFP florescence, (B, F, J) VOZ1-RFP fluorescence and (C, G, K) merged BNP-GFP and VOZ1-RFP fluorescence. (D, I, K) merged green and blue fluorescence images. (D, H, L) Pearson and Spearman´s colocalization test between BNP-GFP and VOZ1-RFP signals. Bars = 10 µm.

**Table S1.** Clones retrieved by a yeast two-hybrid screen using BNP (aa1-250) as a bait.

| Locus | Annotation | # clones | PBS | Length in aa (interaction region) | Localization |
|---|---|---|---|---|---|
| At2g42400 | VOZ2, Vascular plant one zinc finger protein 2 | 4 | D | 450 (1-450) | Cytosol[a] Nucleus[a] |
| At1g28520 | VOZ1, Vascular plant one zinc finger protein 1 | 2 | C | 486 (129-486) | Nucleus, cytosol |
| At2g44370 | BNP, Binucleate pollen | 3 | B | 250 (1-250) | Late endosome/ Prevacuolar Compartment/ Multivesicular bodies[b] |
| At4g32460 | BIIDXI, pectin methylesterase activator | 4 | D | 365 (160-365) | Endoplasmatic reticulum, Golgi |
| At1g10770 | PMEI, pectin methylesterase inhibitor | 1 | D | 167 (1-167) | Plasma membrane[c] |
| At5g20070 | NUDX19, Nudix hydrolase 19 | 55 | A | 438 (60-438) | Chloroplast[d] |
| At1g79690 | NUDT3, Nudix hydrolase 3 | 1 | D | 772 (344-772) | Cytosole, vacuole[f] |
| At5g14660 | PDF1B, peptide deformylase 1B | 30 | A | 273 (1/273) | Chloroplast[g] |
| At1g52410 | TSA1 (TSK-associating protein 1) | 17 | A | 759 (562/759) | Small cytoplasmic vesicles[h] |
| At1g48540 | Outer arm dynein light chain 1 protein | 4 | B | 1063 (718/1063) | Nucleus |
| At4g18740 | Rho termination factor | 2 | C | 245 (1/245) | Nucleus |
| At1g53580 | GLY3, glyoxalase II 3 | 2 | C | 294 (3/294) | Mitochondrion |
| At4g32520 | SHM3, serine hydroxymethyltransferase 3 | 3 | C | 529 (194/529) | Plastid |
| At1g75990 | 26S proteasome regulatory subunit S3 | 1 | D | 487 (256/487) | Cytosol Nucleus |

| | | | | | |
|---|---|---|---|---|---|
| At5g40770 | ATPHB3, Prohibitin 3 | 1 | D | 277 (1/277) | Mitochondrion |
| At5g41790 | CIP1, COP1-interactive protein 1 | 2 | D | 1499 (1416/1499) | Cytoskeleton[i] Cell membrane[j] |
| At2g28640 | EXO70H5, exocyst complex component 7 | 2 | D | 605 (40/605) | Mitochondrion |
| At4g32460 | Protein of unknown function, DUF642 | 4 | D | 365 (160/365) | Extracellular |
| At2g38670 | PECT1, ethanolamine-phosphate cytidylyltransferase | 1 | D | 421 (186/421) | Mitochondrion outer membrane[k] |
| At3g46850 | Subtilase family protein | 2 | D | 529 (194/529) | Extracellular |
| At1g04620 | Coenzyme F420 hydrogenase beta subunit | 1 | D | 462 (6/462) | Chloroplast[l] |
| At3g49960 | Peroxidase | 1 | D | 329 (1/329) | Extracellular |
| At2g41080 | Tetratricopeptide repeat (TPR)-like superfamily protein | 1 | D | 650 (598-650) | Microsome |

[a-l] Indicates experimentally proved; otherwise means predicted. [a](Yasui et al. 2012), [b](present work), [c](Zhang et al. 2010) , [d](Ogawa et al. 2008), [e](Ito et al. 2011), [f] (Carter et al. 2004), [g](Dirk et al. 2001), [h](Suzuki et al. 2005), [l](Ren et al. 2016), [j](Matsui et al. 1995), [k](Mizoi et al. 2006), [l](Meguro et al. 2011). Interaction region is defined as the sequence shared by all prey fragments matching the same reference protein. PBS (Predicted Biological Score) column represents confidence of interaction; A: Very high confidence in the interaction, B: high confidence, C: good confidence, D: moderate confidence.

**Table S2**. Primers used in the study.

| Primer | Sequence (5´ to 3´) |
|---|---|
| *BNP* Promoter+ORF -Fw | CACCGCTTGTGATTGTTCTTTCTTTTG |
| *BNP* Promoter+ORF -Rv | GATCAACTTGAGACAAGCCTTT |
| *BNP* Promoter -Fw | CACCGCTTGTGATTGTTCTTTCTTTTG |
| *BNP* Promoter -Rv | CATGATGGCTAATATCTTCTTCTTCT |
| *BNP* ORF-Fw | CACCATGGCCGCAAGAAAACCGTC |
| *BNP* ORF-Rv | GATCAACTTGAGACAAGCCTTT |
| BIFC-*BNP* ORF-Fw | CACCGCCGCAAGAAAACCGTCGG |
| BIFC- *BNP* ORF-Rv | TTAGATCAACTTGAGACAAGCCTT |
| BIFC-*VOZ1* ORF -Fw | CACCAAGGCTAAGAACCGTGTTGATGA |
| BIFC-*VOZ1* ORF -Rv | TCAGGGGATATAATAGTCGCTTAG |
| BIFC-*VOZ2* ORF -Fw | CACCTCAAACCACCCGAAGATCACATC |
| BIFC-*VOZ2* ORF -Rv | TCACTCCTTACGACCTTTGGTTGG |
| BIFC-*VLG* ORF -Fw | CACCGCTTCCCGCCCTTCAGTGAGAC |
| BIFC-*VLG* ORF -Rv | GTTAATATCTTAGATCATTTTAAGC |
| *VOZ1*-RFP-Fw | CACCAAGGCTAAGAACCGTGTTGATGA |
| *VOZ1*-RFP -Rv | TCAGGGGATATAATAGTCGCTTAG |
| *VOZ2*-RFP -Fw | CACCTCAAACCACCCGAAGATCACATC |
| *VOZ2*-RFP -Rv | TCACTCCTTACGACCTTTGGTTGG |
| *bnp*-1 -LP | ACGGTAGAGCAATGTGAGTGG |
| *bnp*-1 -RP | AGACCCAGCGTAATTGACCTC |
| LBb1 | GCGTGGACCGCTTGCTGCAACT |
| GKGT8474 | ATAATAACGCTGCGGACATCTACATTTT |
| *bnp*-2 -RP | TTGAAAATTTTGTTTTCTCGTGTG |
| *bnp*-2 -LP | GATTATCTGGAACAGTCTCTTGGC |
